# The composition of piRNA clusters in *Drosophila melanogaster* deviates from expectations under the trap model

**DOI:** 10.1101/2023.02.14.528490

**Authors:** Filip Wierzbicki, Robert Kofler

## Abstract

It is widely assumed that the invasion of a transposable element (TE) in mammals and invertebrates is stopped when a copy of the TE jumps into a piRNA cluster (i.e. the trap model). However, recent works, which for example showed that deletion of three major piRNA clusters has no effect on TE activity, cast doubt on the trap model. Therefore, we aim to test the trap model. We show with population genetic simulations that the composition of regions that act as transposon traps (i.e. possible piRNA clusters) ought to deviate from regions that have no effect on TE activity. Next, we investigated TEs in five *D. melanogaster* strains using three complementary approaches to test whether the composition of piRNA clusters matches these expectations. We found that the abundance of TE families inside and outside of piRNA clusters is highly correlated, although this is not expected under the trap model. Furthermore, we found that the distribution of the number of TE insertions in piRNA clusters is also much broader than expected, where some families have zero cluster insertions and others more than 14. One feasible explanation is that insertions in piRNA clusters have little effect on TE activity and that the trap model is therefore incorrect. Alternatively, dispersed piRNA producing TE insertions and temporal as well as spatial heterogeneity of piRNA clusters may explain some of our observations.

## Introduction

Transposable elements (TEs) are short sequences of DNA that selfishly spread in host organisms, even if this selfish activity reduces the fitness of the host [Orgel and Crick, 1980, Doolittle and Sapienza, 1980, Hickey, 1982]. The ability to transpose within the host genome increases the chance of the TE to be transmitted to the next generation [Hickey, 1982]. TEs are highly successful invaders that can be found in virtually all species investigated so far [Wicker et al., 2007]. They show a large diversity in sequence, structure and mechanisms for propagation [Wicker et al., 2007, Sultana et al., 2017].

Although many examples of beneficial TE insertions have been reported [Casacuberta and González, 2013], it is believed that most TE insertions are either neutral or deleterious [Arkhipova, 2018, Pasyukova et al., 2004]. The fact that organisms with highly active TEs are frequently infertile strongly supports the idea of the deleterious effects of TEs [Kidwell et al., 1977, Pélisson, 1979, Moon et al., 2018, Wang et al., 2018, Brennecke et al., 2008]. Additionally, the observation that TEs are rare in coding regions but abundant in non-coding regions is thought to be largely due to negative selection against TEs [Bartolomé et al., 2002, Bergman et al., 2006, Kofler et al., 2012]. Furthermore, the shift of the site-frequency-spectrum of TEs towards rare alleles is frequently interpreted as support for negative selection against TEs [Petrov et al., 2003, Barrón et al., 2014]. The spread of TEs needs to be curbed as an unrestrained accumulation of deleterious TE insertions may drive host populations to extinction [Brookfield and Badge, 1997, Kofler, 2019, 2020]. It was initially believed that TE invasions are solely controlled at the population level by negative selection against TEs [Charlesworth and Charlesworth, 1983, Charlesworth and Langley, 1989]. However, the discovery of small RNA-based host defence mechanisms profoundly changed our view and shifted the attention of many researchers from population genetic control of TE invasions to the functional implications of the host defence. In mammals and invertebrates, the host defence operates via piRNAs, small RNAs ranging in size from 23 to 29 nucleotides [Brennecke et al., 2007, Gunawardane et al., 2007]. These piRNAs mediate the repression of TEs at both the transcriptional and the post-transcriptional level [Brennecke et al., 2007, Gunawardane et al., 2007, Le Thomas et al., 2013, Sienski et al., 2012]. Most piRNAs are produced from distinct source loci termed “piRNA clusters” which in total account for about 3.5% of the *Drosophila* genome [Brennecke et al., 2007]. Two distinct piRNA pathways operate in *Drosophila*, one in the germline and the other in the soma, which mostly controls endogenous retroviruses that invade the germline via virus-like particles produced in the somatic tissue surrounding the germline [Song et al., 1997, Malone et al., 2009]. These two pathways rely on distinct sets of piRNA clusters. The germline pathway is based on dual-strand clusters while the somatic pathway primarily relies upon a single uni-strand cluster, *flamenco* [Malone et al., 2009]. piRNA clusters are largely found in pericentromeric regions [Brennecke et al., 2007, Wierzbicki et al., 2021]. It was later discovered that some TE insertions outside of piRNA clusters are also able to generate piRNAs [Shpiz et al., 2014, Mohn et al., 2014]. Such dispersed piRNA producing source loci (DSL) were found for many different TE families [Shpiz et al., 2014, Mohn et al., 2014]. The mechanism which converts TE insertions into turncoats, which support the host defence rather than the propagation of the TE, is based on maternally transmitted piRNAs [de Vanssay et al., 2012, Le Thomas et al., 2014, Goriaux et al., 2014]. Maternally inherited piRNAs bound to PIWI proteins mediate the installation of chromatin marks at TE insertions that are necessary for piRNA production [Le Thomas et al., 2014, Goriaux et al., 2014]. More recently, it was suggested that siRNAs may also drive the conversion of a TE insertion into a piRNA producing locus [Luo et al., 2022].

Under the current prevailing theory, the trap model, a TE invasion is stopped when a copy of the TE jumps into a piRNA cluster which then triggers the production of piRNAs that silence the TE [Bergman et al., 2006, Malone et al., 2009, Zanni et al., 2013, Goriaux et al., 2014, Yamanaka et al., 2014, Ozata et al., 2019]. Several lines of evidence support the trap model. First, a single insertion in a piRNA cluster, such as X-TAS or 42AB, is able to silence a reporter [Josse et al., 2007, Luo et al., 2022]. Second, an artificial sequence inserted into a piRNA cluster led to the production of piRNAs complementary to the inserted sequence [Muerdter et al., 2012]. Third, deletion of ZAM from the somatic piRNA cluster *flamenco* led to derepression of ZAM. Later the host reacquired the ability to suppress the TE likely due to a ZAM insertion in a germline cluster [Duc et al., 2019, Yoth et al., 2022]. Fourth, computer simulations showed that piRNA clusters are able to stop TE invasions, even in the absence of negative selection against a TE [Kofler, 2019]. Fifth, studies monitoring TE invasion in experimental populations showed that piRNAs complementary to the newly invading TE rapidly emerged and that the generation of piRNAs was accompanied by the emergence of insertions in piRNA clusters [Kofler et al., 2018, 2022]. On the other hand, it was also shown that the observed number of cluster insertions at later generations, where the TE is likely silenced by the host, was lower than expected under the trap model [Kofler et al., 2018, 2022]. Additionally, computer simulations showed that piRNA clusters are solely able to control TE invasions if the clusters have a minimum size (as fraction of the genome) and that these minimum size requirement are barely met in some species [Kofler, 2020]. Finally, deletion of three major piRNA clusters in the germline of *D. melanogaster* did not lead to an activation of TEs [Gebert et al., 2021]. Due to these conflicting results, it is an important open question as to whether the trap model holds. Here, we argue that population genetics can shed light on this issue. Since TEs spread in populations, we argue that a complete understanding of TE invasions requires a synthesis of functional and population genetic considerations. Such a synthesis can lead to surprising outcomes. One notable example comes from the number of cluster insertions necessary to stop a TE invasion. Functional work suggests that a single TE insertion in a piRNA cluster may be sufficient to silence a TE [Josse et al., 2007, Luo et al., 2022]. However, even when assuming that a single insertion is sufficient to stop a TE, population genetic models suggest that at least four insertions per diploid individual are necessary to stop a TE invasion. This can be explained by the fact that most TE insertions in piRNA clusters will be segregating in the population, and that recombination among these segregating cluster insertions will lead to a heterogeneous distribution of cluster insertions in the next generation, where some individuals will carry many cluster insertions and some solely a few or even none [Kofler, 2019]. The TE will be active in the individuals without cluster insertions and thus the average number of cluster insertions in the population will increase. Only when individuals carry around four cluster insertions, do most individuals in the population end up with at least a single insertion. Interestingly, this requirement for four cluster insertions was robust over a wide range of different parameters and scenarios [Kofler, 2019]. This stability in the number of cluster insertions required to silence a TE invasion led us to speculate that the composition of regions that act as transposon traps (e.g. possibly piRNA clusters) should differ markedly from regions that have no effect on TE activity. Such composition differences would provide us with an opportunity to test the trap model. We first performed computer simulations under the trap model and indeed found that the composition of transposon traps should differ from regions having no effect on TE activity in two important aspects. Firstly, for transposon traps, we do not expect a positive correlation between the abundance of TEs within and outside of the trap region, while such a correlation is expected for regions having no effect on TE activity. Secondly, we expect a narrow distribution of the abundance of different TE families in transposon traps, with few families having less than one or more than 14 insertions in transposon traps. By contrast, the expected distribution of TE insertions in regions having no effect on TE activity is much wider. Interestingly, the observed composition of piRNA clusters in five different *D. melanogaster* strains is not in agreement with expectations under the trap model but rather suggests that the composition of piRNA clusters resembles regions having no effect on TE activity. Finally, we suggest amendments to the trap model that may account for the observed discrepancies. In particular, we think that dispersed source loci (DSL) and spatial or temporal heterogeneity of piRNA clusters may account for the observed composition of piRNA clusters.

## Results

It is an important open question whether TE copies inserting into piRNA clusters are responsible for stopping TE invasions (i.e. the trap model). Here, we argue that population genetics can shed light on this issue, as it makes testable predictions about the composition of regions that act as transposon traps (possible piRNA clusters). Notably, the composition of transposon traps should differ markedly from reference regions that have no effect on TE activity.

In this work we proceed in two steps. First, we use simulations to identify key differences in the composition between regions that act as transposon traps (trap model) and reference regions that have no effect on TE activity (random model). Second, we test whether the observed composition of piRNA clusters in five *D. melanogaster* stains best fits with expectations under the trap model or the random model.

### Simulations of TE invasions

In a previous simulation study where we investigated TE invasions under the trap model, we realized that TE invasions are typically controlled when diploid individuals carry, on average, around four insertions in transposon traps (possible piRNA clusters), although we assumed that a single trap insertion per diploid is sufficient to silence the TE [Kofler, 2019]. Recombination and random assortment among segregating trap insertions will lead to a distribution of trap insertions in populations, where some individuals will end up with several trap insertions and others with just a few or even none at all. Only when diploids carry an average of about four trap insertions, will the vast majority of the offspring end up with at least a single trap insertion. The observation that about four trap insertions per diploid individual are necessary to stop TE invasions was highly robust over all evaluated parameters (e.g. different sizes of genomes and piRNA clusters, transposition rates and population sizes; [Kofler, 2019]. Based on this robustness of the number of trap insertions, we hypothesized that the composition of transposon traps should deviate from the composition of reference regions having no effect on TEs in two aspects. First, the distribution of the number of insertions for all different TE families should be very narrow in transposon traps (most families should have around 2 insertions in transposon traps per haploid genome) while the distribution should be much broader for reference regions. Second, for the different TE families, the abundance of TEs in reference regions and the rest of the genome should be highly correlated whereas no correlation is expected between the TE abundance in transposon traps and the rest of the genome. We performed extensive simulations of TE invasions with Invade [Kofler, 2019] to validate our hypotheses about these two key differences in the composition of transposon traps and reference regions. The choice of parameters for the simulations was inspired by *D. melanogaster*. We simulated 5 chromosome arms with a uniform recombination rate of 4cM/Mb. On one end of each chromosome we simulated transposon traps and on the other end reference regions (fig. 1A,B). Both the transposon traps and the reference regions each cover 3.5% of the genome, similar to dual-strand piRNA clusters in the germline of *D. melanogaster* [Brennecke et al., *2007]. We assumed a constant transposition rate u* (i.e. the probability that a single TE copy generates a new copy in the subsequent generation) and novel TE insertions were distributed randomly in the genome. We simulated a population size of *N* = 1000 and non-overlapping generations. To avoid the stochastic early stages of an invasion, where a novel TE is frequently lost by genetic drift [Le Rouzic and Capy, 2005, Kofler, 2019], we triggered each invasion by introducing 1000 TE insertions at random genomic positions (starting frequency *p* = 1*/*2 *\** 1000). For each scenario we simulated 300 replicates. Note that replicates of TE invasions may either be interpreted as invasions of the same TE family in different populations (species) or as invasions of different TE families in the same population. In this work we rely on the second interpretation, which allows us to link our simulation results to the observed abundance of the different TE families in piRNA clusters (see below).

**Figure 1:**
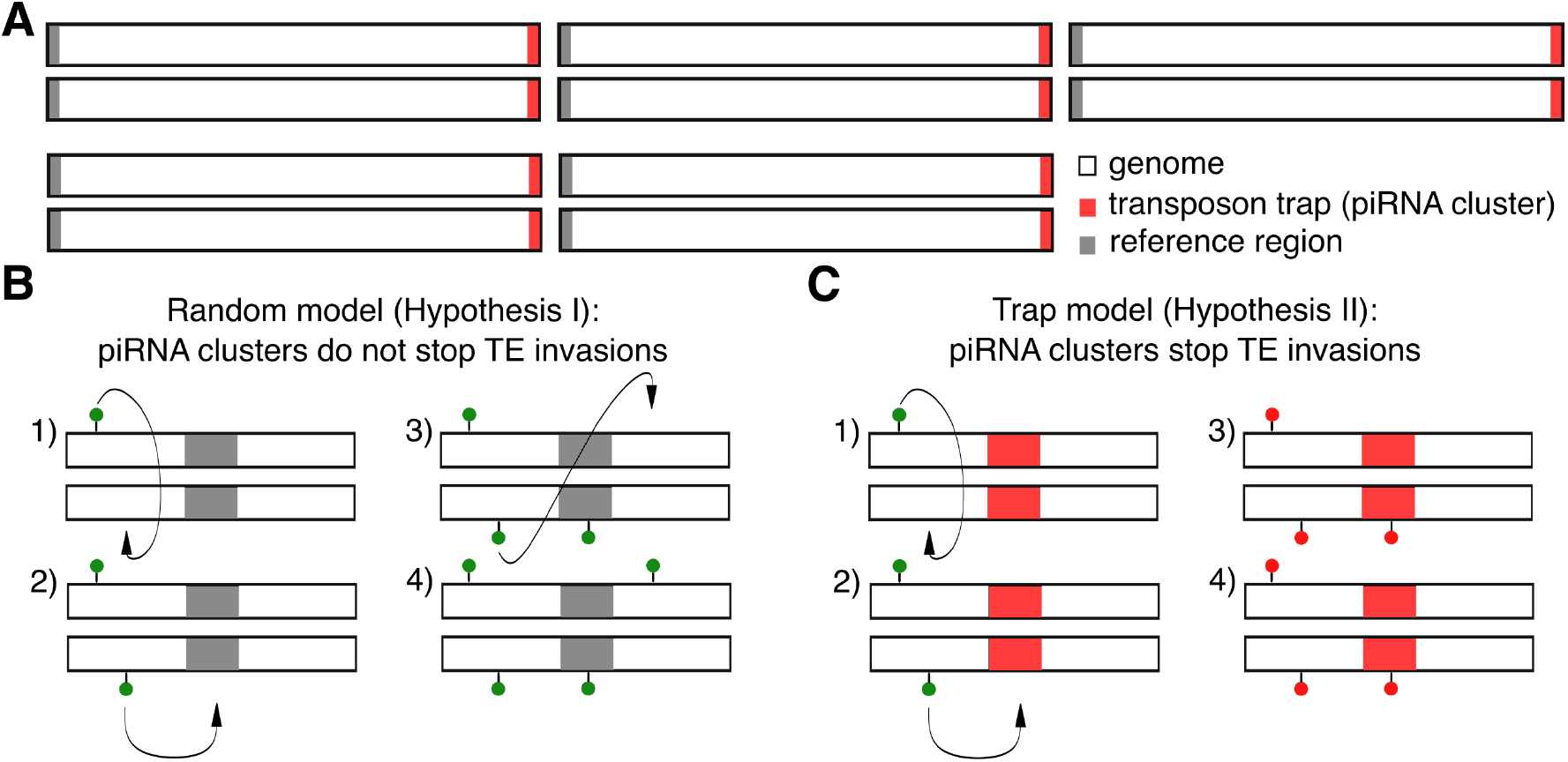
Overview of our simulation approach for testing the trap model. A) We simulated diploid organisms with 5 chromosomes and a uniform recombination rate of 4cM/Mb. Equally sized transposon traps and reference regions were simulated on opposite ends of the chromosomes. We assumed that a TE insertion into a reference region (B) has no effect on TE activity while an insertion into a transposon trap (C) silences the TE. green circle: active TE insertion, red circle: inactive TE insertion

In agreement with recommendations for biological modelling [Otto and Day, 2007], we started with a simple model and than gradually increased the complexity. In the first and simplest scenario, we simulated an identical transposition rate in all replicates (*u* = 0.1) and neutral TE insertions (fig. 2A). At generation 2000 we measured the abundance of TEs in i) the genome, ii) the transposon trap and iii) the reference region. Note that by generation 2000 the invasion is silenced in all replicates, either by segregating or fixed insertions in transposon traps. In a previous study we referred to these distinct stages of TE invasions under the trap model as the shotgun phase and inactive phase, respectively ([Kofler, 2019]; fig. 2, left panels, yellow and red). Even under this simple model we observed a striking heterogeneity in the abundance of TE insertions during the invasion among the 300 replicates (fig. 2A).

**Figure 2:**
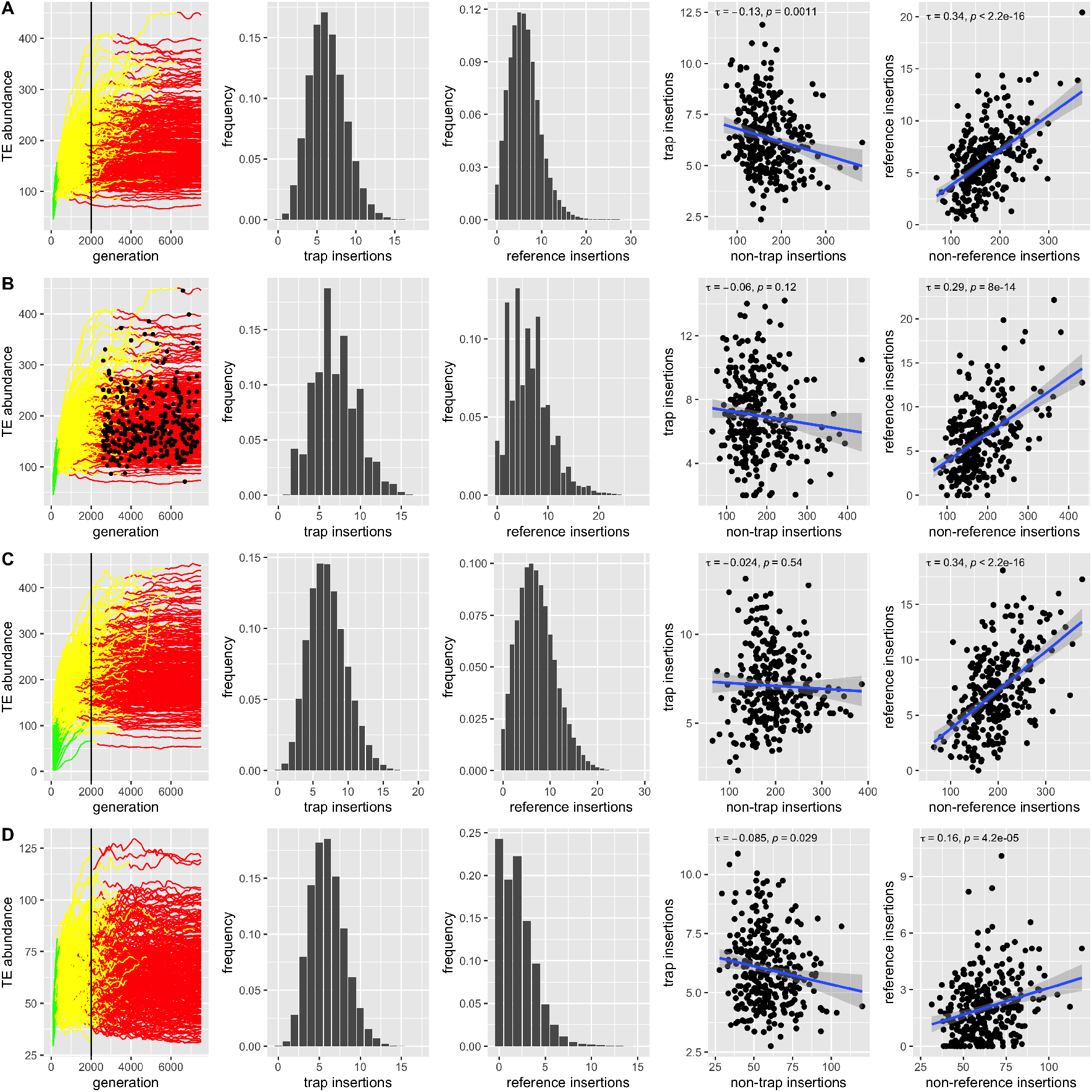
Key differences in the composition of transposon traps (possible piRNA clusters) and reference regions under four different models (A-D). We simulated 300 replicates for each model. From left to right, panels show the TE abundance during invasions, where colors indicate the three distinct phases of TE invasions (invasions controlled by TE insertions in transposons traps are shown in yellow and red; [Kofler, 2019]) (i), a histogram with the abundance of TE insertions in transposon traps (ii), reference regions (iii), and the correlation between the abundance of TE insertions in transposon traps (iv) or reference regions (v) and the rest of the genome. A) A simple model with neutral TE insertions and a constant transposition rate (*u* = 0.1). Invasions were sampled at generation 2000 (black line). B) A simple model with neutral TE insertions and constant transposition rate (*u* = 0.1). Invasions were sampled at different time points between 2500 and 7500 generations (black dots). C) A model with neutral TE insertions. Each replicate has a different, randomly chosen, transposition rate (0.005 *≤ u ≤* 0.5). D) A model where TE insertions have negative fitness effects (site-specific negative effects: 10% of the insertions each have *x* = 0.1, *x* = 0.01, *x* = 0.001 and *x* = 0.0001; 60% are neutral). All replicates have a constant transposition rate (*u* = 0.1).

We first investigated the distribution of the TE abundance in both the transposon traps and reference regions. In agreement with our hypothesis, we found that the distribution of the TE abundance in transposon traps is narrower than that of the reference regions (fig. 2A, second vs third panel). Very few individuals from these 300 replicate populations have less than 1 (0.06%) or more than 14 (0.07%) TE insertions in a transposon trap (fig. 2A, second panel). By contrast, more individuals have less than 1 (2.00%) or more than 14 (1.97%) insertions in reference regions. Importantly, this observation is independent of the size of transposon traps and reference regions (supplementary fig. S1). While it is obvious that individuals with silenced TEs will have at least one insertion in a transposon trap, it is perhaps less clear why more than 14 trap insertions are also not expected under the simple trap model. Since a single TE insertion in a transposon trap silences an invading TE, a continuous accumulation of insertions in the trap regions is not feasible. Only recombination among segregating trap insertions can lead to a slightly elevated number of trap insertions in diploid individuals [Kofler, 2019]. However, the TE distribution resulting from recombination in the transposon traps will be narrower than in reference regions, where in addition to recombination, multiple independent TE insertions may occur (figure 2A).

Second, we investigated the correlation of the TE abundance between transposon traps or reference regions and the rest of the genome. Again in agreement with our hypothesis, we observed a significant positive correlation between the abundance of TEs in reference regions and the rest of the genome but not between transposon traps and the rest of the genome (fig. 2 A; fourth and fifth panel). It is obvious that the abundance of TE families in the genome and a random sample of the genome (i.e. the reference region) will correlate. However, the same relation does not hold for transposon traps, since any TE insertion in the trap will deactivate the TE, thus preventing further accumulation of TEs by transposition. For this reason transposon traps consistently accumulate about 2-3 TE insertions per haploid genome during TE invasions, irrespective of the simulated scenario (different trap sizes, transposition rates, genome sizes, population sizes [Kofler, 2019]).

Next, we aimed to investigate the robustness of those two key differences between transposon traps and reference regions in more complex models. In our simple scenario we assumed that all replicates are sampled at the same time (i.e. 2000 generations after the invasion was triggered). It may, however, be argued that different TE families in an organism (corresponding to replicates in the simulations) are usually captured at different stages of the life cycle of a TE. For example, the P-element invaded *D. melanogaster* populations within the last century, while non-LTR TEs likely invaded thousands of years ago [Schwarz et al., 2021, Kidwell, 1983, Bergman and Bensasson, 2007]. To address this issue, we randomly sampled TE invasions between generation 2500 and 7500 (fig. 2B; black dots in the left panel). We did not sample any invasion at the early stages, where the TE is not yet controlled by insertions in transposon traps (figure 2B; left panel, green; rapid invasion phase [Kofler, 2019]). Our two key differences between transposon traps and reference regions were robust to variation in the sampling time of TE invasions (figure 2B).

So far, we have assumed that all replicates (interpreted as different TE families) have an identical transposition rate. It is, however, likely that different TE families have different transposition rates [Nuzhdin and Mackay, 1995, Émilie Robillard et al., 2016, Kofler et al., 2018]. Although, the influence of the transposition rate on TE copy numbers is likely small under the trap model, variation in the transposition rate could lead to an accumulation of different TE copy numbers during invasions [Kofler et al., 2015, Kofler, 2019, Tomar et al., 2022]. To address this, we randomly selected different transposition rates (between *u* = 0.005 and *u* = 0.5) for each replicate (fig. 2C). Our two key differences between transposon traps and reference regions were robust to variation in the transposition rate (fig. 2C).

So far, we only considered neutral TE insertions, i.e. insertions that have no effect on host fitness. While this may be true for many TEs, it is likely that at least some TE insertions have deleterious effects to the fitness of the host [Arkhipova, 2018, Pasyukova et al., 2004]. Therefore, in the next model we considered negative effects of TE insertions (*x*) using the fitness function *w* = 1 *− nx* (*w* fitness, *n* number of TE insertions). For example, an individual that carries 2 TE insertions with negative effects of *x* = 0.1 has, on average, 20% less offspring than an individual without any TE insertions. Initially, we simulated a scenario where all TE insertions have an equal constant effect. To avoid an unlikely equilibrium state between transposition, selection, and piRNA clusters (TSC balance [Kofler, 2019]), we assumed that TE insertions in piRNA clusters are neutral. We performed 300 simulations for each of the following negative effects: *x* = 0.0001, *x* = 0.001 and *x* = 0.01 (supplementary fig. S2A-C). As expected, with weak negative effects (*x* = 0.0001; supplementary fig. S2A), the invasions resemble the neutral scenario (fig. 2A; *Nx <* 1). Large negative effects (*x* = 0.001, *x* = 0.01) had a notable impact on the abundance of TEs during the invasions (supplementary fig. S2B,C). While negative selection had a minimal effect on the abundance of TEs in trap regions, the abundance in reference regions was markedly reduced, with many individuals having zero insertions in reference regions (supplementary fig. S2B,C). Furthermore, we found a positive correlation between the TE abundance in the genome and reference regions but not with transposon traps (where the correlation was actually negative; supplementary fig. S2). Even when we further relax our assumptions by considering negative selection against all TE insertions, including insertions in transposon traps, our two key differences are robust (supplementary fig. S3).

Thus far, we have assumed that all TE insertions have an equal negative effect on host fitness, irrespective of the genomic insertion site. It is, however, possible that different TE insertions have diverse fitness effects depending on the insertion site ([Pasyukova et al., 2004, Houle and Nuzhdin, 2004]). For example, insertions into coding sequences are likely more harmful than insertions in intergenic regions. We evaluated the effect of heterogeneous fitness effects of TE insertions using the linear fitness function: *w* = 1 *−∑ n*_*i*_ where *n*_*i*_ is the negative effect of each TE insertion. With such a site-specific model we may vary i) the effect size of the TE insertions and ii) the proportion of the genome at which a TE insertion will lead to the given negative effects. We varied the fraction of sites where a TE insertions causes negative fitness effects (*x* = 0.01) from 10% to 70%. The remaining 90% to 30% of the possible insertion sites were neutral. Our two key differences between trap and reference regions were robust to variation in the number of neutral insertion sites (supplementary fig. S4). Next, we considered a more complex distribution of site-specific deleterious effects of TE insertions. We assumed that 10% of the insertions each have a negative of *x* = 0.1, *x* = 0.01, *x* = 0.001 and *x* = 0.0001, while the remaining 60% of the sites were neutral (fig. 2 D). Our key differences were again robust (fig. 2 D).

Up to this point, we have simulated TE insertions with identical fitness effects among replicates (corresponding to TE families). It could, however, be argued that different TE families have diverse fitness effects. For example, a TE family with an insertion bias into promotor regions may be, on average, more deleterious than a TE family that has an insertion preference for intergenic regions. In agreement with this, previous work suggests that negative effects of TE insertions may vary among TE families [Charlesworth and Langley, 1989, Barrón et al., 2014]. To consider such a scenario, we performed simulations with different negative effects. For each replicate, we randomly picked a different negative effect between *x* = 0.001 and *x* = 0.1 for 40% of the sites in the genome while insertions into the remaining 60% were neutral. Within a replicate, all non-neutral TE insertions had the same negative effect (supplementary fig. S5). Under this scenario, we once again found a narrow distribution of the TE abundance within traps and no correlation of the TE abundance between trap regions and the rest of the genome (supplementary fig. S5).

In summary, we identified two key differences in the composition of transposon traps and reference regions that were robust in all evaluated scenarios. If piRNA clusters act as transposon traps, we expect i) no positive correlation between the abundance of TEs in piRNA clusters and the rest of the genome and ii) a narrow distribution of the abundance of TE insertions in piRNA clusters, with few individuals having less than 1 or more than 14 cluster insertions for a given TE family.

### The TE composition of piRNA clusters is not in agreement with expectations under the trap model

We next asked whether the observed composition of piRNA clusters is more in agreement with expectations under the trap or the random model, based on the two key differences between transposon traps and reference regions identified above. To address this question, we investigated the TE composition in five *D. melanogaster* strains (Canton-S, DGRP-732, Iso-1, Oregon-R and Pi2). For all five strains, Illumina paired-end reads and genome assemblies are publicly available ([The modENCODE Consortium et al., 2010, Mackay et al., 2012, Hoskins et al., 2015, Chakraborty et al., 2019, Ellison and Cao, 2020, Wierzbicki et al., 2022]). We ensured that the assemblies are of high quality, having complete assemblies of most piRNA clusters (supplementary table S1). Based on the number of completely assembled piRNA clusters, the five assemblies analyzed in this work are among the best out of 37 high-quality (mostly based on long reads) assemblies of diverse *D. melanogaster* strains [Rech et al., 2022] (supplementary table S2). Ovarian small RNA data are available for Canton-S, DGRP-732, Iso-1, Oregon-R ([Song et al., 2014, Fast et al., 2017, Schwarz et al., 2021]). For this work, we generated ovarian small RNA data for Pi2. We employed three complementary approaches to estimate the composition of piRNA clusters in these five strains. First, we identified TE insertions in piRNA clusters based on paired-end reads aligned to the *D. melanogaster* reference genome (release 5; [Hoskins et al., 2007]). Based on the standard annotations of piRNA clusters [Brennecke et al., 2007] and our tool PoPoolationTE2, which locates TE insertions in a reference genome using paired-end data, TE insertions in piRNA clusters and the rest of the genome were identified [Kofler et al., 2016]. This approach has the advantage that it relies on the widely-used standard annotations of piRNA clusters but the disadvantage that some TE insertions in piRNA clusters may be missed. Since piRNA clusters are evolving rapidly [Gebert et al., 2021, Wierzbicki et al., 2021], the sequences of piRNA clusters in the five strains may have diverged from the reference genome. Therefore, some strain-specific reads may not align to the reference genome and thus not all TE insertions may have be identified. Second, we identified TE insertions in the assemblies of the investigated strains using RepeatMasker [Smit et al., 2013-2015] and performed a lift-over of the annotations of piRNA clusters from the reference genome to the assemblies with our CUSCO approach, where the positions of piRNA clusters are identified using unique sequences flanking the reference clusters [Wierzbicki et al., 2022]. In addition to clusters being flanked by two unique sequences, we also included telomeric piRNA clusters into the analysis (e.g. X-TAS) [Josse et al., 2007, Asif-Laidin et al., 2017]. This approach has the advantage that the composition of the rapidly evolving piRNA clusters may be more accurately captured as we rely on assemblies of the investigated strains. Furthermore, this approach is based on the widely used standard annotations of piRNA clusters in *D. melanogaster* [Brennecke et al., 2007]. The disadvantage is that the positions of some reference clusters cannot be identified in the assemblies (for some reference clusters unique flanking sequences could not be identified and some flanking sequences cannot be mapped to the assemblies). Finally, it is possible that, in addition to the composition of piRNA clusters, the location of piRNA clusters is also evolving rapidly [Gebert et al., 2021]. To address this issue, we employed a third approach, where we performed a *de novo* annotation of piRNA clusters in the assemblies of the five strains. We used strain-specific small RNA data for the annotation of piRNA clusters. TE insertions were again identified with RepeatMasker [Smit et al., 2013-2015]. This approach has the benefit that strain-specific variation in both, the location and the composition of piRNA clusters is taken into account, but it has the downside that the locations of these clusters have not yet been substantiated by complementary approaches. For example, apart from an enrichment of piRNAs, dual-strand clusters of the germline typically also show an enrichment of H3K9me3 methylation marks and of Rhino and Kipferl binding sites [Mohn et al., 2014, Baumgartner et al., 2022]. This information is not yet available for the *de novo* annotated piRNA clusters. For all three complementary approaches, we only considered TE families being active in the germline.

We first examined the correlation between the abundance of different TE families in piRNA clusters and the rest of the genome (fig. 3). Under the trap model, we expect either no correlation or a negative correlation (figs. 2, 3A) whereas under the random model a positive correlation is expected (see above). We found a positive correlation between the average number of TE insertions in piRNA clusters and the rest of the genome, with all three approaches for quantifying the TE abundance in the five *D. melanogaster* strains (fig. 3B,C,D). This correlation can also be found if each strain is analyzed separately (supplementary fig. S6).

**Figure 3:**
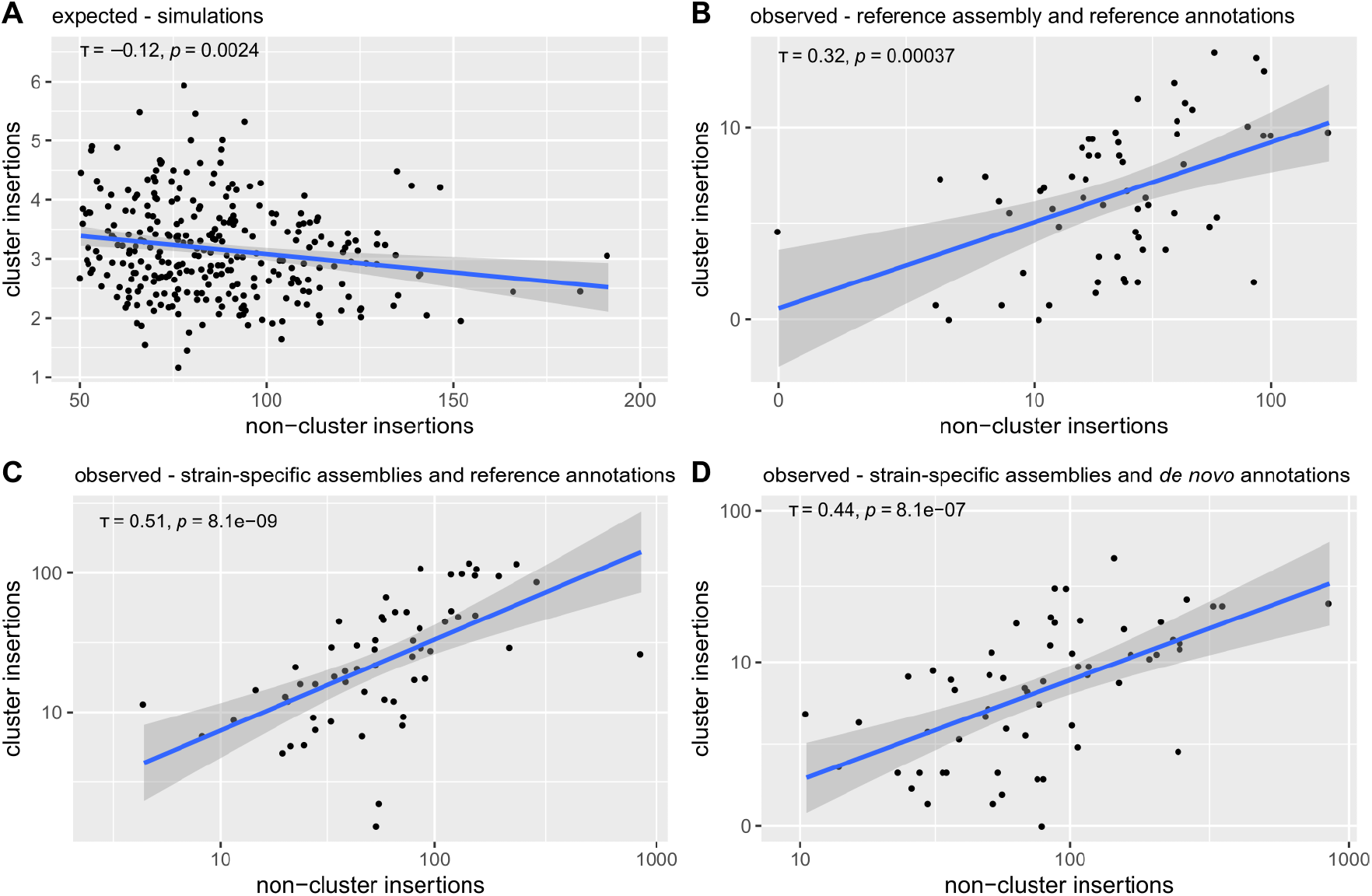
Expected and observed correlation between the TE abundance in piRNA clusters and the rest of the genome. A) Expected correlation based on simulations under the trap model (neutral insertions and *u* = 0.1). B) Observed correlation based on short reads aligned to the reference genome and the reference annotations of piRNA clusters. C) Observed correlation based on strain-specific assemblies and a lift-over of the reference annotations of piRNA clusters using unique sequences flanking the clusters. D) Observed correlations based on strain-specific assemblies and *de novo* annotations of piRNA clusters. All counts refer to copy numbers per haploid genome. For the observed data we averaged the counts over the five strains. Solely TE families active in the germline were considered. Kendall rank correlation coefficients are reported.

In both assembly-based approaches, the TE copy numbers are estimated with RepeatMasker, which occasionally provides fragmented annotations for highly-diverged TE insertions or TEs with internal deletions. It may be argued that such fragmented TE insertions could lead to wrong correlations between the TE content in clusters and the rest of the genome. To address this issue, we repeated the analysis using two different approaches. We first merged fragmented TE annotations with the tool Onecodetofindthemall.pl [Bailly-Bechet et al., 2014] and again found a correlation between the TE abundance within and outside of piRNA clusters (supplementary fig. S7A; based on the assemblies and reference clusters). Second, we used a highly conservative approach solely considering contiguous full-length insertions and again found the correlation between the TE abundance within and outside of piRNA clusters (supplementary fig. S8A; based on the assemblies and reference clusters). Finally, gaps in the assemblies of piRNA clusters indicate assembly problems [Wierzbicki et al., 2022]. Therefore, we repeated the analysis by excluding clusters with gaps but again found a significant correlation between the TE abundance within and outside of piRNA clusters (supplementary fig. S9A).

Next, we focused on the abundance of the different TE families in piRNA clusters. Our simulations show that the abundance of TE families in transposon traps should follow a narrow distribution, with no family having less than 1 and only a few having more than 14 insertions (fig. 2, 4A). Based on our three complementary approaches, we estimated the abundance of each TE family in the haploid genomes of the five strains. We found that the observed distribution of the TE abundance in piRNA clusters differs substantially from expectations under the trap model (fig. 4). First, several TE families do not have a single cluster insertion (fig. 4B,C,D). For example, we could not find cluster insertions for the R2-element, Tirant, Bari1, flea and jockey in some strains (supplementary table S3) Second, many families have many more insertions in piRNA clusters than expected (fig. 4B,C,D).

**Figure 4:**
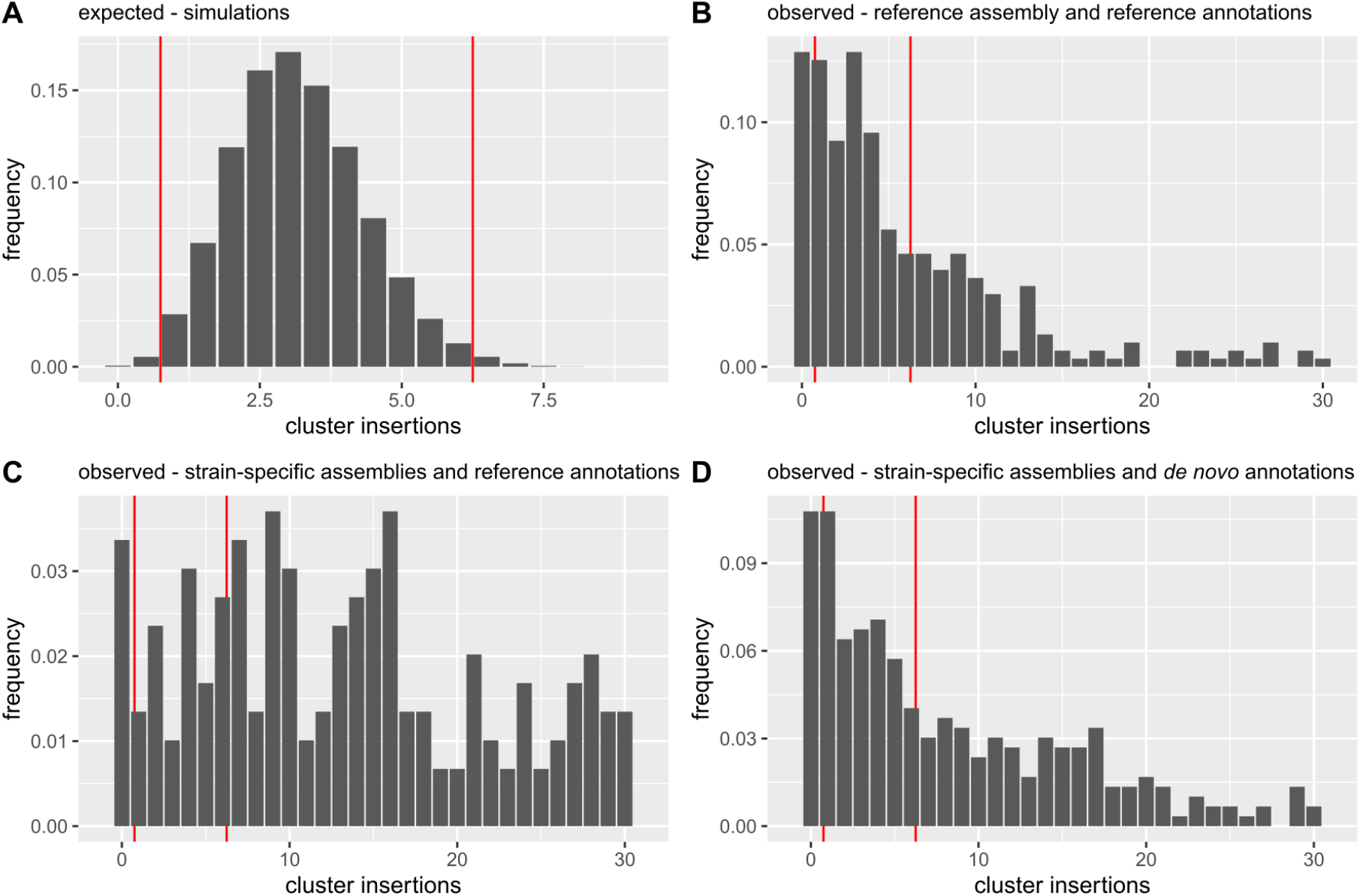
Expected and observed abundance of different TE families in piRNA clusters. A) Histogram showing the expected abundance based on simulations under the trap model (for haploid genomes). B) Observed abundance based on short reads aligned to the reference genome and the reference annotations of piRNA clusters. C) Observed abundance based on strain-specific assemblies and a lift-over of the reference annotations of piRNA clusters, using unique sequences flanking the clusters. D) Observed abundance based on strain-specific assemblies and *de novo* annotations of piRNA clusters. All counts refer to copy numbers per haploid genomes. For the observed data, we averaged the abundance in the five investigated strains. Only TE families active in the germline were considered. At least 98% of the simulations under the trap model are between the red lines. The x-axis was truncated at 30 insertions.

As mentioned above, the assembly-based approaches rely on RepeatMasker, which occasionally provides fragmented annotations for TE insertions. Such fragmented annotations could boost the number of TE insertions in piRNA clusters causing the observed over-representation of some TE families in piRNA clusters. To address this issue, we repeated the analysis by merging fragmented annotations with the tool Onecodetofindthemall.pl [Bailly-Bechet et al., 2014] and again found an over-representation of several TE families in piRNA clusters (supplementary fig. S7B). This over-representation of TEs in piRNA clusters is also found when we just consider full-length insertions of TEs or piRNA clusters assembled without gaps (supplementary figs. S8B, S9B). This analysis also revealed that many TE families (27.3%) do not have a single full-length insertion in a piRNA cluster (supplementary fig. S8B).

To summarize, we find a correlation between the abundance of TEs in piRNA clusters and the rest of the genome, contrary to expectations under the trap model. Moreover, we observed that many TE families either do not have a single insertion in a piRNA cluster or have a larger than expected number of cluster insertions. These observations are more consistent with the random model, which posits that TE insertions in piRNA clusters have no effect on TE activity.

### Abundance of dispersed piRNA producing TE insertions

For several TE families we did not find a single cluster insertion, which is unexpected if piRNA clusters control TE invasions. Apart from assembly problems (see Discussion), there is an alternative hypothesis which may account for the missing cluster insertions. Recently, Gebert et al. [2021] showed that three major piRNA clusters can be deleted with no effect on the activity of the TEs. They suggest that this is due to redundancy in the host defence, where dispersed TE insertions may also produce piRNAs. These dispersed source loci (DSL) could compensate for the missing cluster insertions. TE families without cluster insertions should thus have at least one DSL. Such DSL have a distinct piRNA signature that can be recognized in the genome. piRNA production frequently extends from the TE into the genomic regions flanking the TE insertion, such that antisense piRNAs are produced upstream of the TE and sense piRNAs downstream of the TE [Shpiz et al., 2014]. To identify DSL, we scanned the assemblies of the five strains for TE insertions flanked by these asymmetric piRNA signatures (for example fig. 5A). To evaluate the performance of our algorithm for finding DSL, we computed the fraction of conserved genes (BUSCO genes) with asymmetric piRNA signature. Since none (or almost none) of the conserved BUSCO genes are expected to act as a piRNA producing locus, this approach provides us with an estimate of the fraction of false positive DSLs identified by our approach. We found very few BUSCO genes with such asymmetric piRNA signatures and therefore argue that our approach has a high specificity (fig. 5B). Using our approach, we estimate that about 2 *−* 5% of the TE insertions are piRNA source loci outside of piRNA clusters (fig. 5B). The abundance of DSL varies among the TE families and the strains (supplementary fig. S10). DSL were more evenly distributed along chromosomes than insertions in piRNA clusters, which were most abundant near centromeres (supplementary fig. S11). Finally, we asked whether the DSL could compensate for the missing TE insertions in piRNA clusters. Indeed, we found a DSL for most of the TE families without a single cluster insertion (fig. 5 C, supplementary fig. S12, supplementary table S3; based on the strain-specific assemblies and the reference clusters).

**Figure 5:**
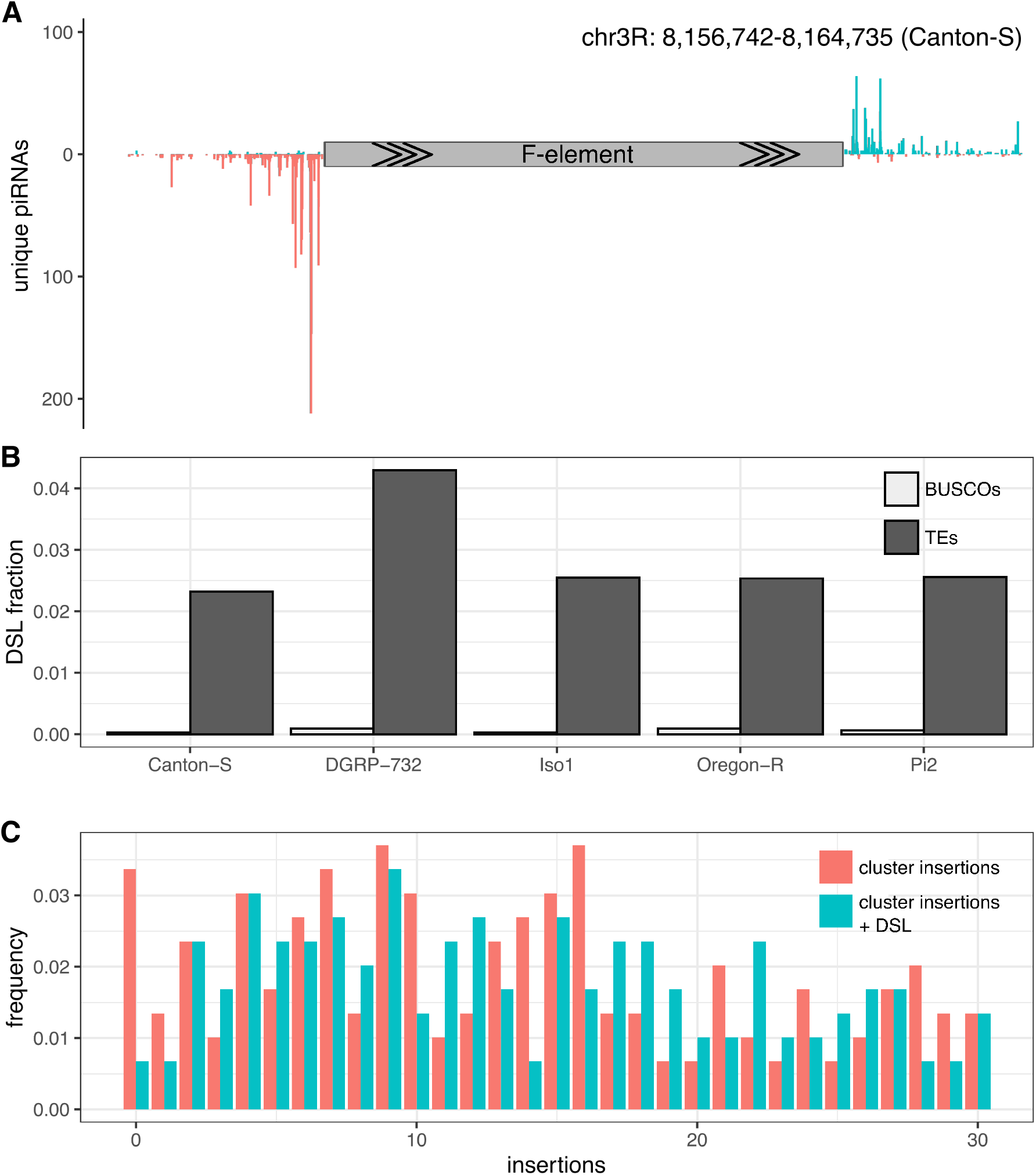
Dispersed piRNA producing source loci (DSL) could compensate for the missing cluster insertions. A) Example of a DSL in Canton-S. Note the typical signature of DSL where antisense piRNAs (red, negative y-axis) align upstream of the TE insertions (F-element) and sense piRNAs (blue, positive y-axis) downstream of the TE [Shpiz et al., 2014]. B) Abundance of DSL in the five investigated strains (dark grey). As a negative control we also computed the fraction of BUSCO genes with a typical DSL signature (light grey; BUSCO genes should not have DSL signatures). C) Abundance of piRNA producing loci in the five strains. Data are shown for cluster insertions (red) as well as cluster insertions plus DSL (cyan). Note that the number of families without piRNA producing locus is dramatically reduced when DSL are considered.

Only Tirant and Bari1 in Oregon-R do not have a single piRNA producing locus (neither cluster insertion nor DSL; supplementary table S3).

In summary, our results show that we find at least one piRNA producing locus, either a cluster insertion or a DSL, for most of the recently active TEs in *D. melanogaster*. Therefore, DSL can largely compensate for the missing cluster insertions of some TE families.

## Discussion

In this work, we showed that the observed composition of piRNA clusters in five *D. melanogaster* strains is not in agreement with expectations under the simple trap model, i.e. the notion that a single insertion in a piRNA cluster stops the proliferation of TEs.

Based on extensive simulations of TE invasions under the trap model, we identified two key differences between genomic regions where a TE insertion represses TE activity and regions where insertions have no effect on TE activity (i.e. transposon traps vs reference regions). First, the abundance of different TE families should be more narrowly distributed in transposon traps than in reference regions. Second, the abundance of TEs within and outside of reference regions should be positively correlated, whereas no positive correlation is expected for transposon traps. These differences are robust over a wide range of different scenarios and parameters, such as varying transposition rates, sizes of piRNA clusters, age of the invasions, negative effects of TEs, and different combinations of these factors. However, we can not fully rule out the possibility that some specific parameter combination or simulation scenario exists where the two key differences do not hold. With simulations, only a finite number of possible scenarios or parameter combinations can reasonably be explored. Nevertheless, our work shows that under the vast majority of the feasible scenarios, for example, a positive correlation between the TE abundance within and outside of clusters is not expected and that a parsimonious conclusion is that piRNA clusters (or large portions of piRNA clusters; see below) do not act as transposon traps. When simulating the TE composition of reference regions, we assumed that some other region outside of the reference act as transposon traps. It is however possible that TE invasions are not stopped by transposon traps but by an hitherto unknown mechanism (possibly siRNAs; see below [Luo et al., 2022]). However, even in such a scenario the observed correlation of the TE abundance between reference regions and the rest of the genome will persist. Additionally, the distribution of TE insertion in reference regions will likely become even more heterogeneous than observed in our simulations, where we assumed that some region with the same size as the reference region acts as transposon trap. Therefore, we think that our two key differences between reference regions and transposon traps are conservative.

We examined the observed distribution of piRNA clusters in five *D. melanogaster* strains. We restricted the analyses to these strains because high-quality genome assemblies, genomic reads and small RNA data from ovaries are only available for these strains (small RNA data for Pi2 were generated by us). In principle, a single strain would have been sufficient to test our predictions about the composition of piRNA clusters under the trap model but an analysis of five strains provides a more comprehensive picture, allowing us to rule out that our results are merely based on a strain that may, for example, have assembly problems. Furthermore, we tested the observed TE composition using 3 complementary approaches, each with their own strengths and weaknesses.

Our analysis of the five strains revealed that not a single cluster insertion can be found for several TE families, which is in stark contrast to expectations under the trap model. We can not fully rule out the hypothesis that some insertions in the piRNA clusters were missed, since the assemblies of the five strains may still be incomplete. Only telomere-to-telomere assemblies of the investigated strains, currently available for a single human genome [Nurk et al., 2022], will provide a complete picture of the genomic landscape of *Drosophila*, including its piRNA clusters. However, we consider it unlikely that the missing cluster insertions are a result of insufficient assembly quality. First, apart from the reference genome (Iso-1), the assemblies used in this work are based on long reads, which enable high-quality assemblies even for highly repetitive regions [Khost et al., 2017, Chakraborty et al., 2018, Ellison and Cao, 2020, Wierzbicki et al., 2022]. In agreement with this, multiple quality metrics suggest that the assemblies of the five strains used in this work are of high quality (supplementary table S1). Furthermore, we found the location of the unique sequences flanking the piRNA clusters of the reference genome in most of our assemblies (91.8-97.6%). In comparison with other recently published long-read assemblies [Rech et al., 2022], our assemblies are among those with the most completely assembled piRNA clusters (supplementary table S2). Finally, we confirmed that the number of cluster insertions is insufficient for many TE families with an approach that does not rely on the assemblies of the individual strains, but instead is based on short reads aligned to the reference genome. Taken together, we do not think that an insufficient assembly quality can account for the missing cluster insertions. One other possible hypothesis which could explain the missing cluster insertions is that some of the TE families may not yet be silenced by the piRNA pathway. For example, a TE family that is currently spreading in *D. melanogaster* could simply not yet have acquired any insertions in piRNA clusters. Previous genomic scans showed that four TE families invaded *D. melanogaster* during the last century: P-element, I-element, Tirant and hobo [Schwarz et al., 2021]. Since several of our strains were sampled early during the last century, not all of these four TEs are present in our five strains (e.g. the P-element is missing in Iso-1). Therefore, we solely considered TE families that were actually present in a given strain for the analysis of missing cluster insertions (for example we did not consider the P-element in Iso-1). However, Tirant is present in the analyzed Oregon-R assembly [Chakraborty et al., 2019] but we did not find a cluster insertion. It is thus feasible that Tirant is not yet under host control in this line.

We believe that the most likely explanation for the missing cluster insertions is that euchromatic TE insertions, outside of piRNA clusters, could also be producing piRNAs (DSL: dispersed piRNA producing source loci [Shpiz et al., 2014]). The conversion of regular TE insertion into a DSL may be driven by maternally deposited piRNAs [de Vanssay et al., 2012, Hermant et al., 2015, Le Thomas et al., 2014]. Hence, once the piRNAs targeting an invading TE have emerged, large numbers of TE insertions outside of piRNA clusters may be converted into piRNA producing loci. The DSL generate a substantial redundancy in the number of piRNA producing loci and could thus compensate for the missing cluster insertions. In agreement with this, a recent study demonstrated that the deletion of three major piRNA clusters had no effect on the activity of the resident TE families [Gebert et al., 2021]. The authors suggested that DSL may compensate for the deleted cluster insertions [Gebert et al., 2021]. Based on our genome-wide scan of the five strains, we suggest that about 2 *−* 5% of all TE insertions act as DSL (supplementary fig. S10). When we consider DSL in addition to cluster insertions, we found that most of the TE families, in all five strains, have at least one piRNA producing locus (either DSL or cluster insertion). We therefore conclude that DSL could account for the missing cluster insertions.

However, the DSL cannot account for the over-representation of some TEs in piRNA clusters, nor the correlation between the TE abundance inside and outside of piRNA clusters. Both the over-representation and the correlation of the TE abundance persisted when we merged fragmented TE insertions, only considered full-length insertions or removed piRNA clusters with assembly gaps from the analysis (supplementary figs. S7, S8, S9). One possible explanation contributing to the over-representation of some families is likely repeat expansion. Hence, some TEs in piRNA clusters may not represent independent insertion events but rather tandem duplications of sub-sequences of the clusters [Wierzbicki et al., 2022]. Both the over-representation of some families and the correlation of the TE abundance are solely expected for random genomic regions, where a TE insertion has no effect on the activity of the TE. Therefore, one possible explanation of our data is that the trap model, which posits that insertions in piRNA clusters stop TE invasion, is incorrect. It is, for example, feasible that other mechansims, such as siRNAs generated from dsRNA of TEs, are responsible for activating the host defence against an invading TE [Luo et al., 2022]. However, we do not consider it likely that the trap model is entirely incorrect, since there is strong evidence that insertions in piRNA clusters can produce piRNAs [Muerdter et al., 2012] Furthermore insertions in the germline clusters X-TAS and 42AB were shown to silence reporter constructs [Josse et al., 2007, Luo et al., 2022]. Additionally, the transposon ZAM was activated in some strain due to loss of a ZAM insertion in the somatic piRNA cluster *flamenco* and silenced at later generations due to a novel insertions in a germline piRNA clusters [Duc et al., 2019, Yoth et al., 2022].

One possible explanation for the over-representation and the correlation is that the number of cluster insertions required for silencing a TE varies among the TE families. It could, for example, be speculated that silencing of short TEs requires more cluster insertions than silencing of long TEs, since short TEs may generate fewer piRNAs. However, so far no evidence exists for such heterogeneity between the TE families.

One potential alternative explanation for both the over-representation and the correlation, is that not the entire sequence of the piRNA clusters acts as random region (where insertions have no effect on TE activity) but rather only certain regions within the clusters. Hence, TE insertions in some clusters may activate the host defence against an invading TE while insertions in other clusters may have no effect. It is even feasible that this silencing capacity varies within a cluster. In agreement with this, previous studies found that the number of TE insertions in X-TAS (a piRNA cluster) necessary to silence a TE varies among strains: in one strain a single insertion was sufficient, while in another strain two insertions were required [Marin et al., 2000, Ronsseray et al., 1996]. This heterogeneity among the strains may be due to different insertion sites of the TE in X-TAS. To test the hypothesis that the silencing capacity varies among (within) clusters, it would be important to insert artificial sequences into many regions of different piRNA clusters and test if these insertions repress a reporter (e.g.: similarly to Luo et al. [2022] and Josse et al. [2007]).

The over-representation and the correlation could also be due to a rapid turnover of the location of piRNA clusters. It is thought that the position of piRNA clusters in the next generation is not determined at the genomic level, e.g. due to sequence motifs, but rather by maternally deposited piRNAs. It is not clear how stably the piRNA composition is inherited over many generations but it is feasible that such epigenetic transmitted information may be subject to some variation over the course of time. In agreement with this, recent studies found a rapid turnover in the location and composition of piRNA clusters [Gebert et al., 2021, Wierzbicki et al., 2021]. This raises the possibility that some of the piRNA clusters were solely established after an invading TE was silenced by the host. For TEs that invaded before a cluster was established, the region of the soon to be established cluster would act much like a random region with no effect on TE activity.

In summary, we think that the heterogeneity of piRNA clusters both temporally (rapid turnover of the location) and spatially (varying silencing capacity within cluster) is likely responsible for both the observed over-representation of some TEs families in piRNA clusters and the correlation of the TE abundance within and outside of piRNA clusters. It may be a promising avenue of future work to investigate this heterogeneity of the clusters.

Our work also raises the important open question as to which role piRNA clusters play in stopping TE activity. It is feasible that piRNA clusters are important for activating the piRNA-based host defence but once the host defence is established, piRNA clusters become dispensible due to the redundancy of piRNA producing loci for example generated by DSL [Chen and Aravin, 2021]. It is also possible that the silencing of an invading TE is not triggered by insertions into piRNA clusters but rather by siRNAs [Luo et al., 2022]. In this scenario, piRNA clusters may be dispensable for both triggering and maintaining the piRNA-based host defence against an invading TE. Replicated TE invasions in strains with a defective siRNA pathway and strains lacking major piRNA clusters may be a promising approach to address these open questions.

## Materials and Methods

### Data of fly strains

In this work we analyzed the five *D. melanogaster* strains Canton-S, DGRRP-732, Iso-1, Oregon-R and Pi2. As a part of our previous work, we generated high-quality assemblies for Canton-S (GCA 015832445.1) and Pi2 (GCA 015852585.1) [Wierzbicki et al., 2022]. We also used the assemblies of the *D. melanogaster* reference strain Iso-1 (release 6; FlyBase [Hoskins et al., 2015], DGRP-732(GCA 004798075.2) [Ellison and Cao, 2020]and Oregon-R (GCA 003402015.1) [Chakraborty et al., 2019]. Genomic short-read data for these strains have been made available (SRR11460805, SRR11460802,SRR11460799,SRR933591,SRX671607) [The modENCODE Consortium et al., 2010, Mackay et al., 2012, Wierzbicki et al., 2022]. We previously published the small RNA data from ovaries of Canton-S (SRR11846567) and Iso-1 (SRR11846566) [Schwarz et al., 2021]. We also used the ovarian small RNA data of DGRP-732 (SRR1572814) [Song et al., 2014] and Oregon-R (SRR4473614) [Fast et al., 2017].

### Small RNA sequencing

To obtain small RNA data for Pi2 we extracted total RNA from ovaries of this strain using TRIzol (Invitrogen, Carlsbad, CA). The small RNA library preparation and sequencing was performed by Fasteris (Geneva, Switzerland). RNAs were separated in a polyacrylamide gel electrophoresis and the abundant 2S rRNA was depleted. The libraries were prepared using the the Illumina TruSeq small RNA kit and sequenced on the Illumina HiSeq systems.

### Simulations

TE invasions were simulated using our tool “Invade” (v0.808, [Kofler, 2019]). For this work, we added a novel feature which allowed us to specify the position of reference regions (i.e. genomic regions with no effect on TE activity). Similar to our previous work [Kofler, 2020], we simulated diploid organism with 5 chromosomes, each 10Mb in size. We used a uniform recombination rate of 4cM/Mb. Transposons traps (piRNA clusters) and reference regions, each accounting for 3.5% of the genome, were simulated on opposite ends of the chromosomes. We assumed that a TE insertion in a transposon trap silences the TE while an insertion into a reference region has no effect. We simulated populations of 1, 000 diploid individuals using non-overlapping generations for 10, 000 generations. To avoid the early stochastic phases of TE invasions, where TEs are frequently lost due to genetic drift [Le Rouzic and Capy, 2005], we triggered each invasion by randomly distributing 1, 000 TE insertions in the population (population frequency *p* = 1*/*2 *\** 1000). Unless mentioned otherwise, we used a constant transposition rate of *u* = 0.1. Individuals with a TE insertion in a piRNA cluster had a transposition rate of *u* = 0.0. Simulations with negative selection were performed using the fitness function *w* = 1 *−∑* ^*n*^ *x*_*i*_ where *w* is the fitness of an individual and *x*_*i*_ the negative effect of each TE insertion. We terminated a simulation when the average fitness fell below *<* 0.1 (extinction of the population). Since Invade reports TE insertions per diploid individuals, TE counts were divided by 2.0 to obtain estimates for haploid genomes.

### Identification of TEs

For the identification of TEs, we used the consensus sequences of TEs in *D. melanogaster* and added the sequences of Chimpo, Chouto, Pifo, Batumi, and Bica [Quesneville et al., 2005] (see data availability). To detect TE insertions based on short-read data, we relied on our tool PopoolationTE2 (v1.10.04) [Kofler et al., 2016]. We use the release 5 of *D. melanogaster* reference genome, as a widely used standard annotation of piRNA clusters is available for this release [Brennecke et al., 2007]. In agreement with the manual of PoPoolationTE2, we first built a FASTA-file consisting of the repeat-masked (RepeatMasker version 4.0.7, [Smit et al., 2013-2015]) reference genome and the consensus sequences of TEs. We mapped the reads to this FASTA-file using bwa mem (version 0.7.17-r1188) and the option -M (mark secondary alignments [Li and Durbin, 2009]. We generated a ppileup file (–map-qual 15), identified signatures of TE insertions (– min-count 2 –signature-window minimumSampleMedian), estimated population frequencies, and paired-up signatures of TE insertions. To exclude unreliable and somatic insertions, we solely considered TEs with a minimum population frequency of 0.3. TE insertions with frequencies lower than 0.6 were assumed to be heterozygous. To obtain the number of TE insertions per haploid genome, the abundance of heterozygous insertions were divided by two (the number of homozygous insertions *≥* 0.6 were not altered). We used RepeatMasker (4.0.7) to identify TE insertions in the assemblies of the five strains (-s -no is -nolow [Smit et al., 2013-2015]). To prevent fragmented TE annotations, we set the *−− frag* option to 40, 000, 000, which is higher than the largest scaffold in our data. We filtered for TE insertions with a minimum length of 100bp and a maximum divergence of 10% to the consensus sequence. Finally, we excluded families that were not recently active (*≥* 25% population frequency [Kofler et al., 2015] and families which are only active in the soma (as these TEs are controlled by a distinct piRNA cluster). The TE families considered in this work are shown in the supplementary table S4.

### Annotation of piRNA clusters

We used our CUSCO approach [Wierzbicki et al., 2022] to lift-over the classic annotation of piRNA clusters to the assemblies of the five strains. We identified sequences flanking the piRNA clusters in (release 5 [Brennecke et al., 2007]) and aligned these sequences to the five assemblies using bwa sw(version 0.7.17-r1188, [Li and Durbin, 2009]). The regions between these two aligned sequences were annotated as piRNA clusters. Telomeric associated sequences (TAS) frequently act as piRNA clusters [Josse et al., 2007] but these clusters are not flanked by a unique pair of sequences. To identify these clusters, we aligned the most distal gene of each chromosome arm (release 6, [Hoskins et al., 2015]) to the assemblies of the five strains and annotated the region between this gene and the end of the contig as TAS cluster. Overlapping piRNA clusters were merged using bedtools (v2.27.1, [Quinlan and Hall, 2010]).

We used proTRAC (V.2.4.4, [Rosenkranz and Zischler, 2012]) and ovarian small RNA data to *de novo* annotate piRNA clusters in the five assemblies. We trimmed reads using cutadapt [Martin, 2011], filtered reads with a length of 23-29nt, and mapped the reads to a set of *D. melanogaster* mRNAs, miRNAs, rRNAs, snRNAs, snoRNAs, tRNAs and TEs [Dos Santos et al., 2015, Quesneville et al., 2005] using NovoAlign (V3.09.00, [Novocraft, 2014]). Based on these alignments, we removed reads mapping to a miRNA, mRNA, rRNA, snRNA, tRNA and snoRNA. Following the proTRAC pipeline, we collapsed overlapping reads, removed low complexity reads, aligned the remaining reads to the assemblies of the five strains, and run pro-TRAC using uniquely mapping reads (-pdens 0.05 -pimin 23 -pimax 29 -1Tor10A 0.3 -clsize 5000 -clstrand 0.5) [Rosenkranz and Zischler, 2012]. Following Gebert et al. [2021], neighboring clusters were joined if the distance between clusters was smaller than their combined lengths.

### Quality of the assembies

We estimated the BUSCO (Benchmarking Universal Single-Copy Orthologs (v5.0.0 [Manni et al., 2021]) for all assemblies using the augustus mode and the *diptera odb10* data set. CUSCO values (the fraction of completely assembled piRNA clusters) were computed based on alignments of unique sequences flanking piRNA clusters (see above) in the reference assembly [Wierzbicki et al., 2022] using bwasw (0.7.17-r1188 [Li and Durbin, 2009]). We delineated between gapped (g.CUSCO) and ungapped CUSCO (u.CUSCO) which only considers piRNA clusters without any assembly gaps. To calculate CQ (coverage quality) and ScQ (softclip quality) values for clusters [Wierzbicki et al., 2022], we mapped long reads of the corresponding strain to the assemblies using minimap2 (v2.16-r922 [Li, 2018]). Only reads with a minimum mapping quality of 60 and minimum read length of 5kb were considered. Assembly sizes and N50 values were obtained from the FASTA index files generated by samtools (v1.9 [Li et al., 2009]).

### DSL detection

For finding the asymmetric piRNA signatures of DSL we solely considered reads with a length between 23 and 29nt. We furthermore excluded reads mapping to miRNAs, rRNAs, snRNAs, snoRNAs and tRNAs. The remaining reads were mapped to the corresponding assembly using NovoAlign (V3.09.00, [Novocraft, 2014]). DSLs were identified based on uniquely mapping reads (minimum mapping quality 5). To estimate the rate of false positive DSL, we considered BUSCO genes with a minimum length of 100bp. For each feature (TE or BUSCO gene), we estimated the number reads aligning 500bp downstream or upstream of the insertion. As DSL we only considered insertions with a minimum of 5 antisense reads per million in the upstream region and 5 sense reads per million in the downstream region. TE insertions in piRNA clusters were not considered as DSL.

## Data availability

The ovarian small RNA data of the *D. melanogaster* strain Pi2 have been deposited in NCBI (BioProject: PRJNA930650). The new version of Invade is available at SourceForge (https://sourceforge.net/projects/invade/; last access on 2/2/2023). All analyses performed in this work were documented with R Markdown and made available at GitHub https://github.com/filwierz/trapmodel (last access on 2/2/2023). The used scripts and the TE library are also available at this repository.

## Author contributions

RK and FW conceived the work. FW extracted RNA, performed simulations and analyzed the data. RK and FW wrote the paper.

## Acknowledgements

We thank all members of the Institute of Population Genetics for feedback and support. We thank Matthew Beaumont for comments. This work was supported by the Austrian Science Fund (FWF) grants P34965-B25 to RK and W1225.

