## supplement for "The composition of piRNA clusters in *Drosophila melanogaster* deviates from expectations under the trap model"

### Supplementary figures

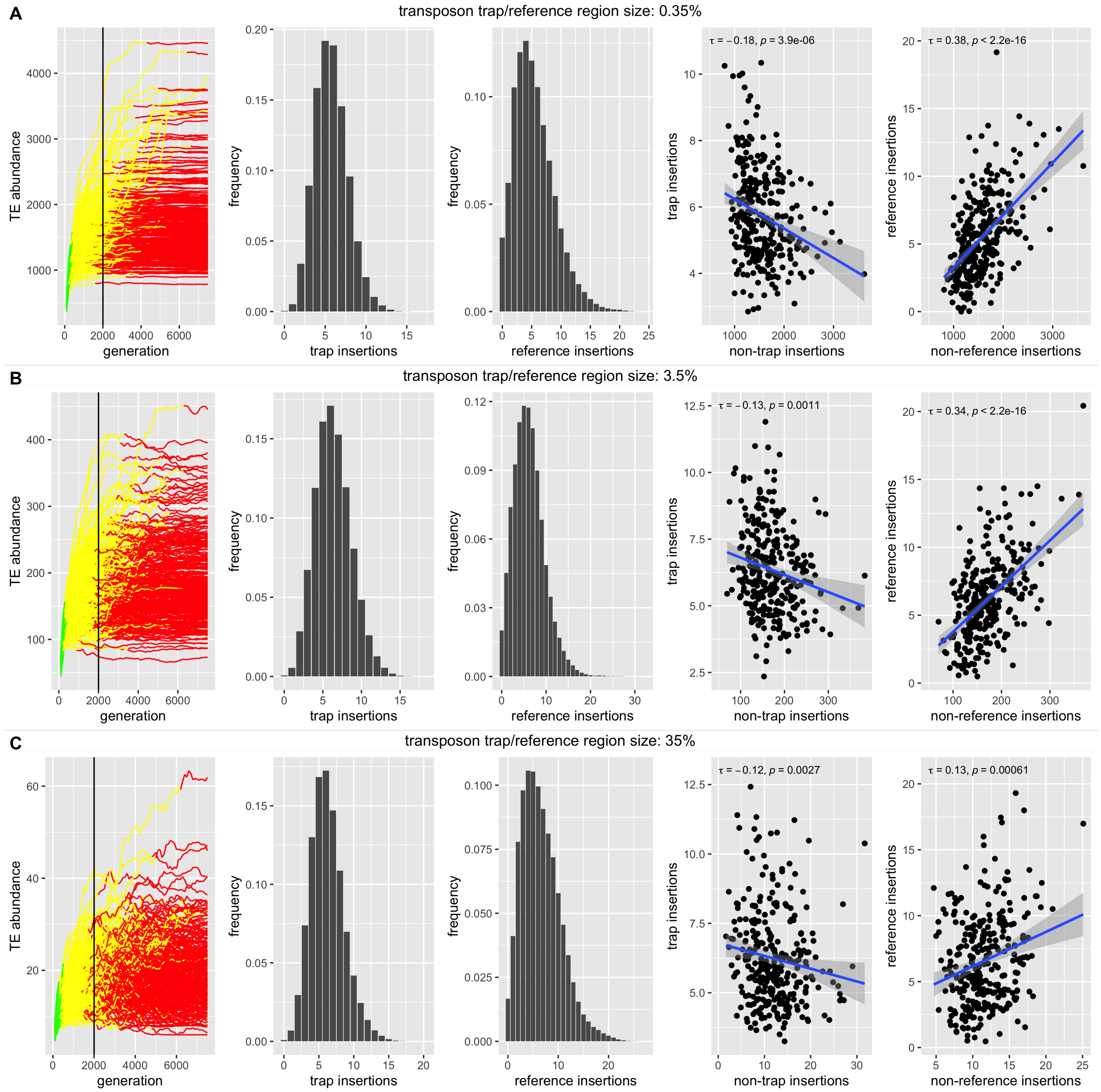

Figure S1: Effect of the size of transposon trap and reference regions on our two key metrics (the distribution and correlation of TEs in trap and reference regions). Neutral simulations with transposon trap and reference region size of A) 0.35% B) 3.5% and C) 35% of the genome. The transposition rate was $u = 0.1$.

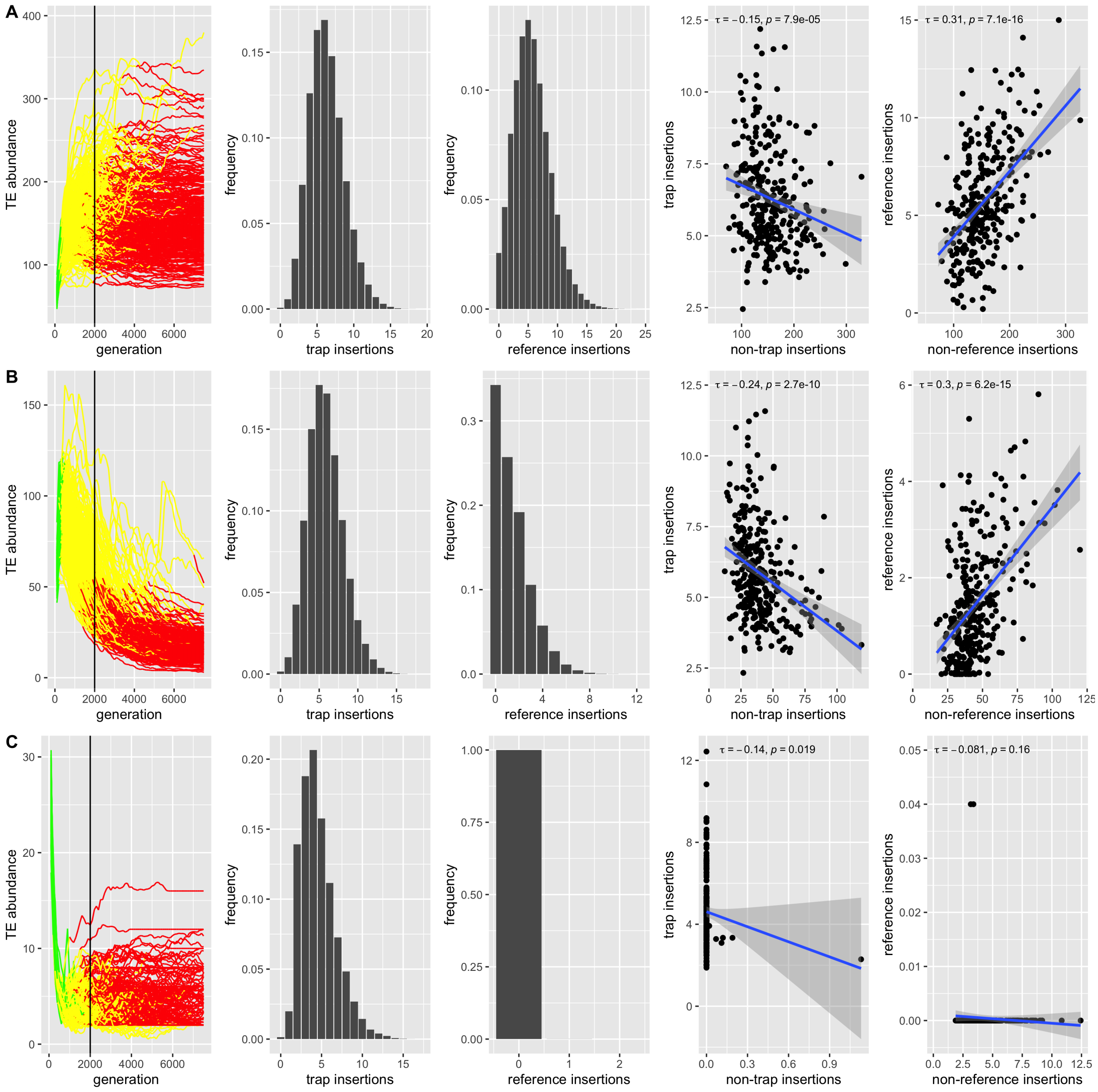

Figure S2: Effect of negative selection, excluding trap insertions, on our two key metrics. Except for TE insertions in transposon traps, each TE insertion reduces the fitness of the host by A) 0.0001 B) 0.001 and C) 0.01. The transposition rate was $u = 0.1$

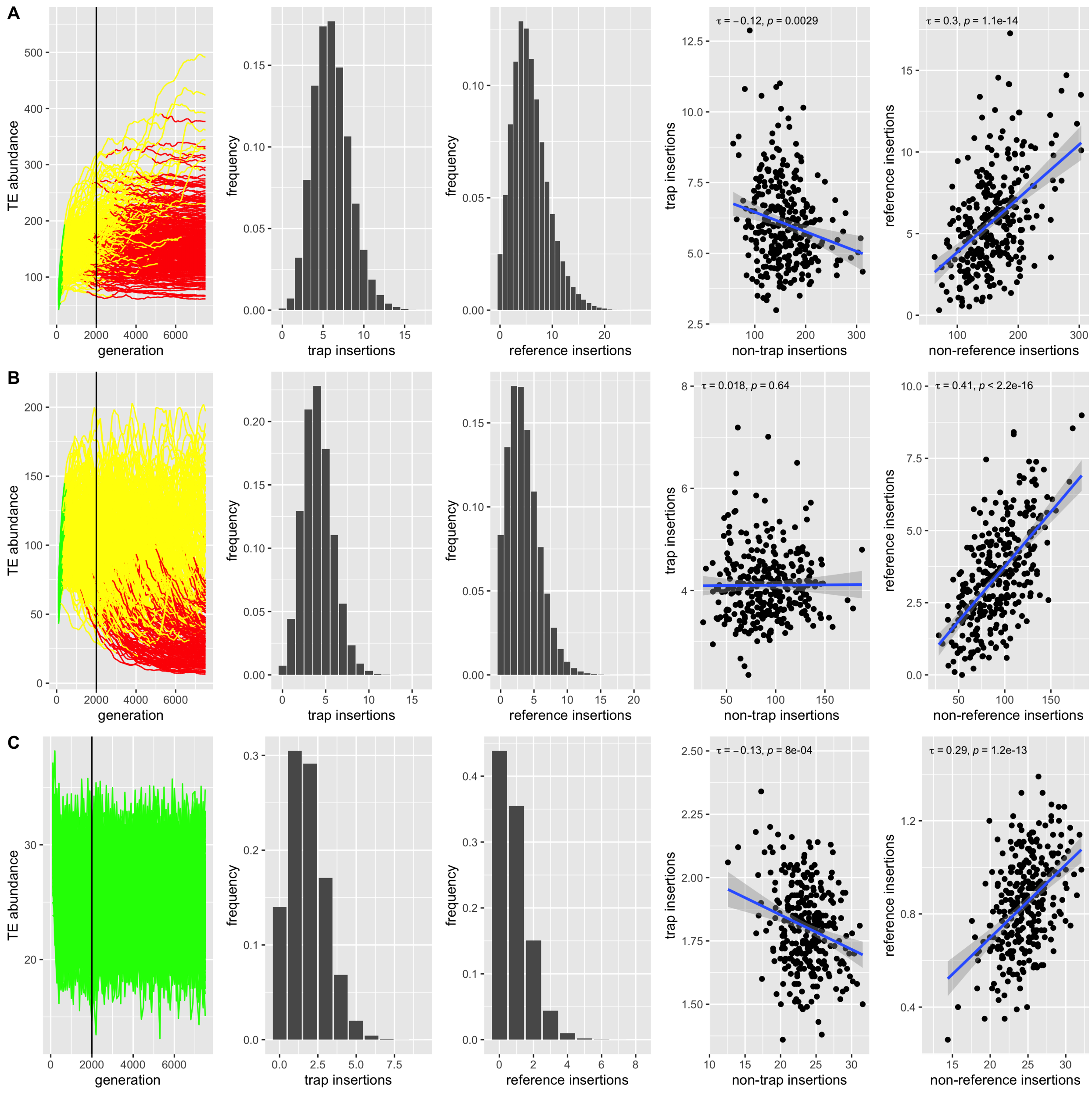

Figure S3: Effect of negative selection, including trap insertion, on our two key metrics. Each TE reduces the fitness of the host by A) 0.0001 B) 0.001 and C) 0.01. The transposition rate was $u = 0.1$. Note that the invasion in C has reached TSC-balance and is thus neither stopped by segregating nor fixed insertions in transposon traps [Kofler, 2019].

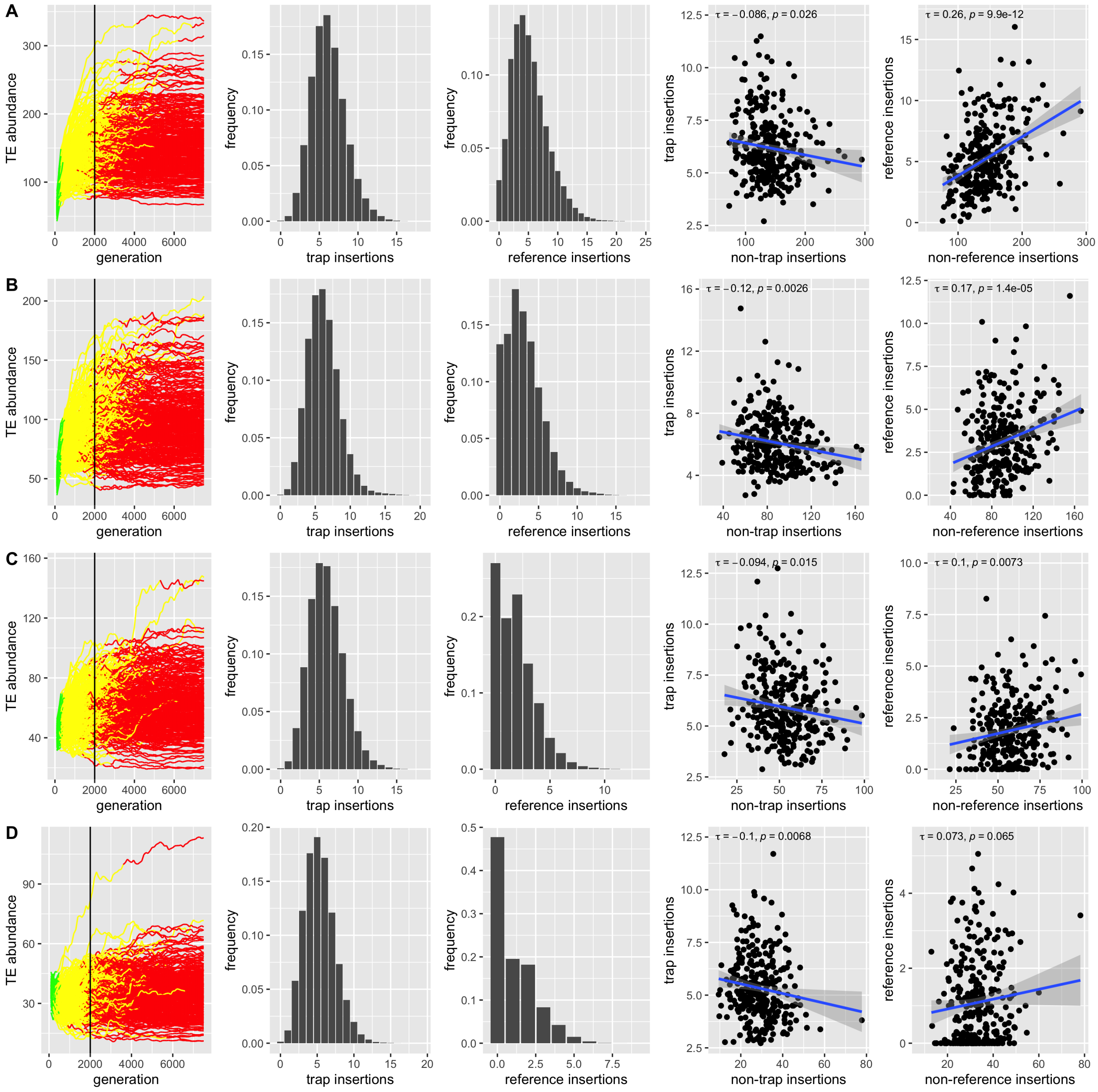

Figure S4: Effect of the fraction of negatively selected TEs on our two key metrics. Except for transposon traps, the following fraction of TE insertions had a negative effect ($x = 0.01$) on host fitness host A) 10% B) 30% C) 50% and D) 70% of the genome. The remaining TE insertions (e.g. for A: $100 - 10 = 90\%$) were neutral ($x = 0.0$). We used a transposition rate of $u = 0.1$ and site-specific negative effects of TEs (hence independent TE insertions at the same site will always have the same fitness effect).

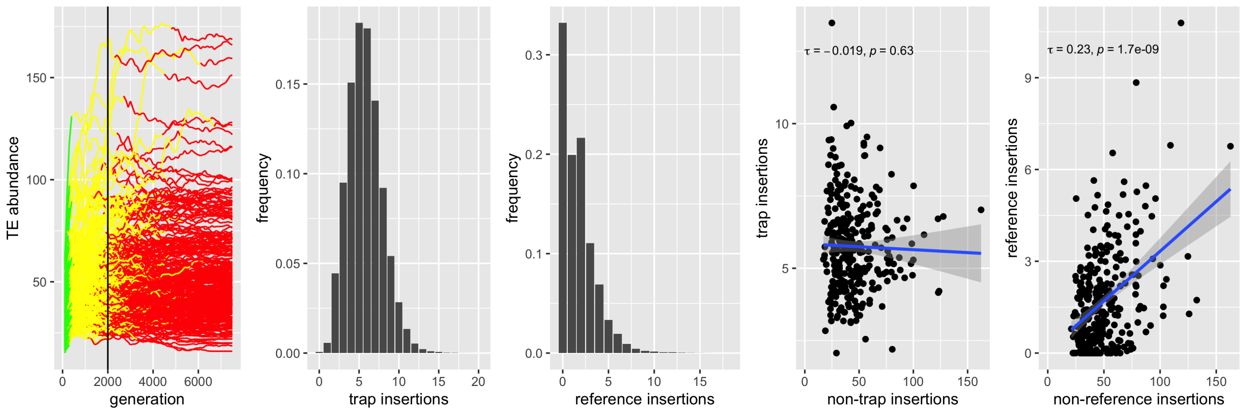

Figure S5: Effect of varying the deleterious effect of TE insertions among replicates on our two key metrics. Each replicate may be regarded as the invasion of a different TE family with distinct fitness effects to hosts. For each of the 300 replicates, we randomly picked a negative effect of TEs between $x = 0.001$ and $x = 0.1$ for 40% of the genome while TE insertions into the remaining 60% were neutral. The transposition rate was $u = 0.1$.

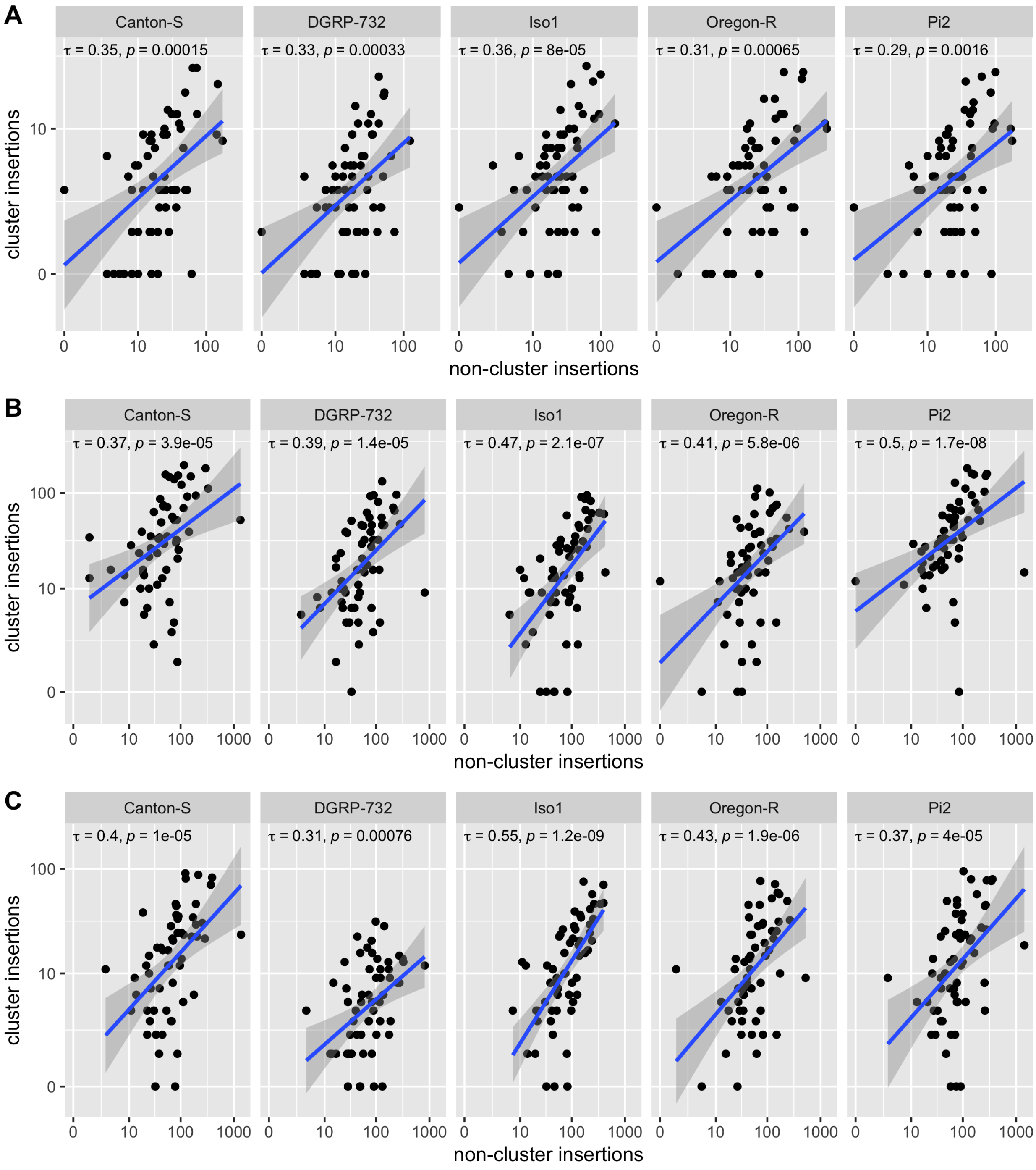

Figure S6: Correlation between the number of TE insertions in piRNA clusters and the rest of the genome (non-cluster insertions). The TE abundance in these two regions was estimated with three different approaches. A) Short reads aligned to the reference genome with the annotations of piRNA clusters of Brennecke et al. [2007]. B) Strain-specific assemblies - annotations of piRNA clusters were lifted-over to the assemblies based on unique sequences flanking the clusters. C) Strain-specific assemblies - *de novo* annotations of piRNA clusters. All counts are copy numbers per haploid genome and Kendall's rank correlations were used.

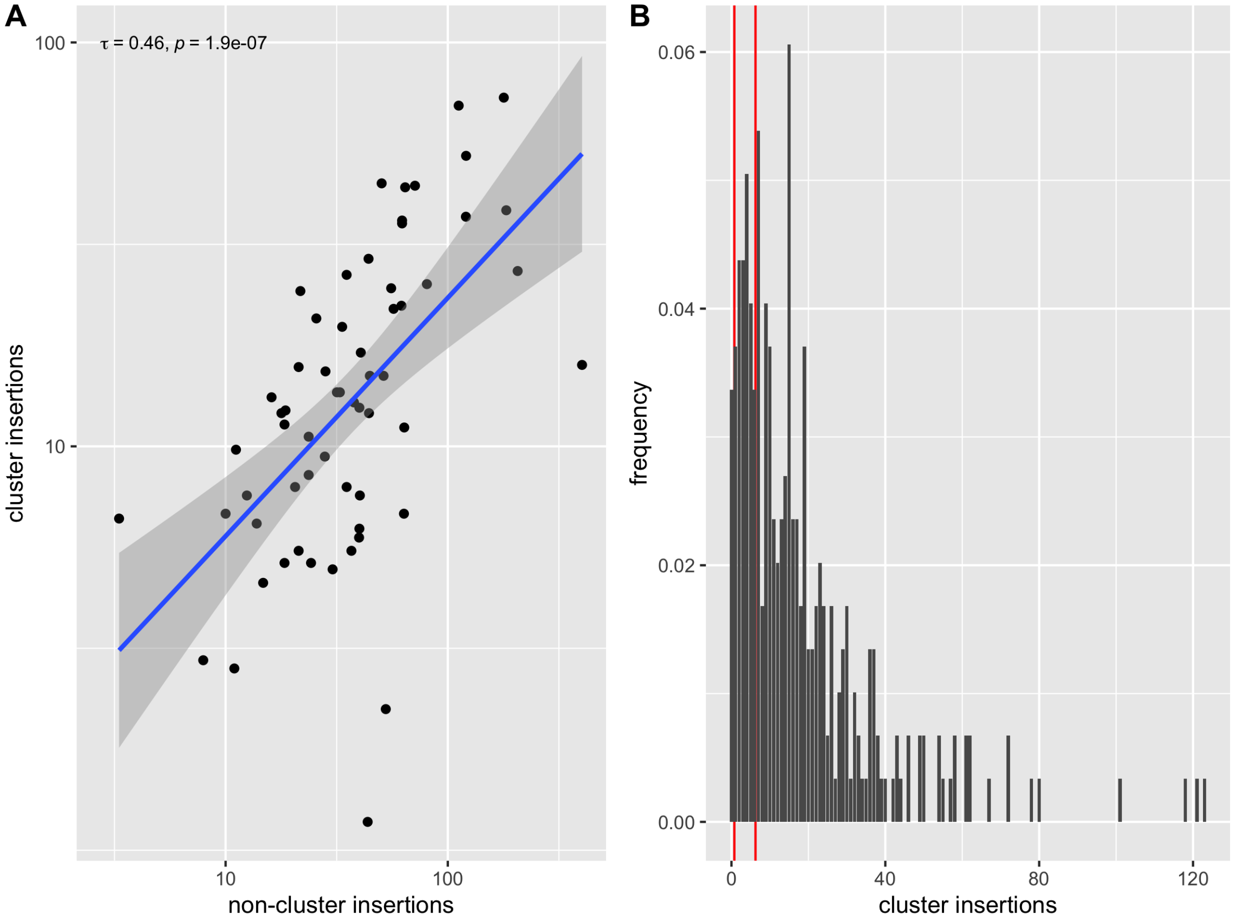

Figure S7: Effect of merging fragmented TEs on the correlation between cluster and non-cluster insertions (A) and the distribution of TE insertions in piRNA clusters (B). Copy numbers are average number of insertions per TE family and haploid genome.

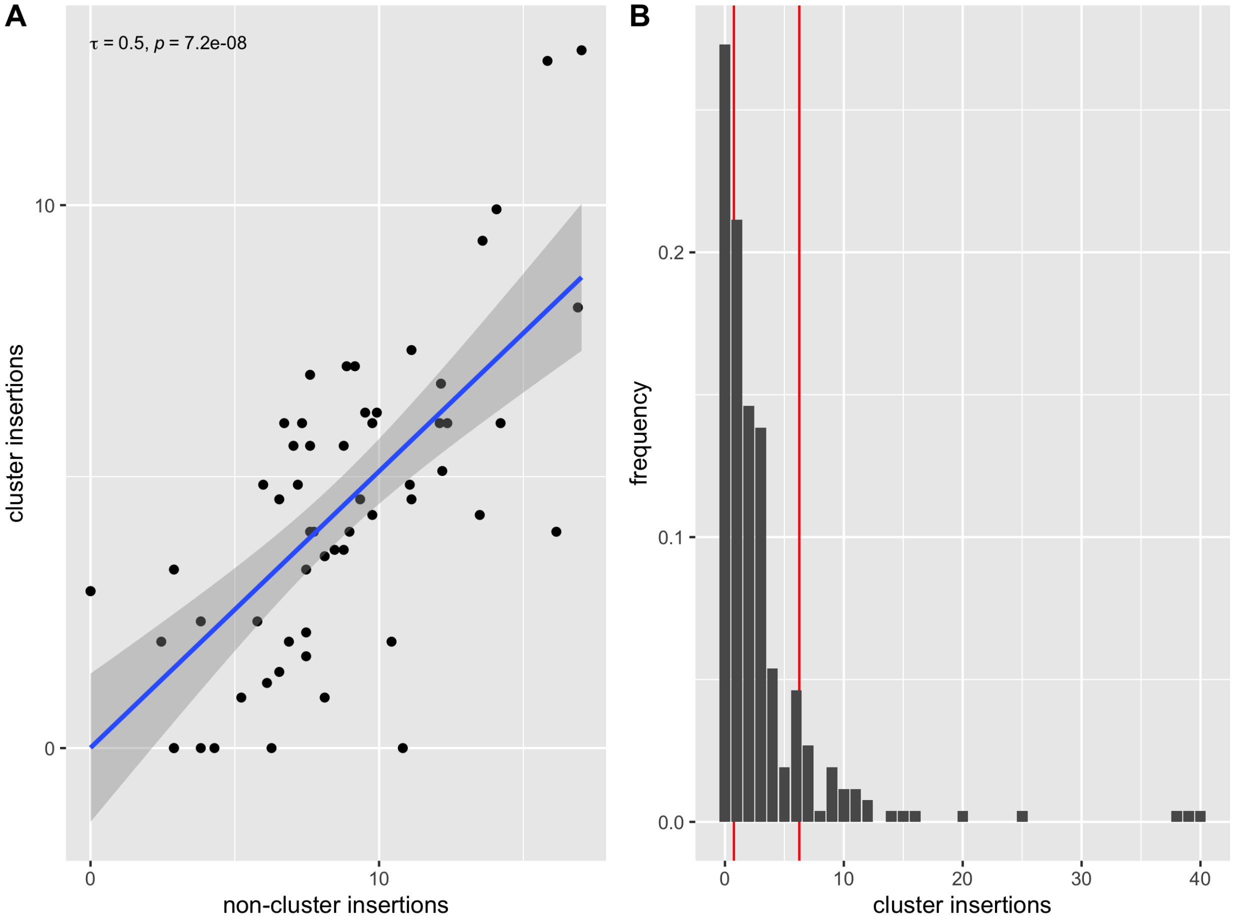

Figure S8: Correlation between cluster and non-cluster insertions (A) and the distribution of TE insertions in piRNA clusters (B) when solely full-length insertions are considered. Copy numbers are average number of insertions per TE family and haploid genome.

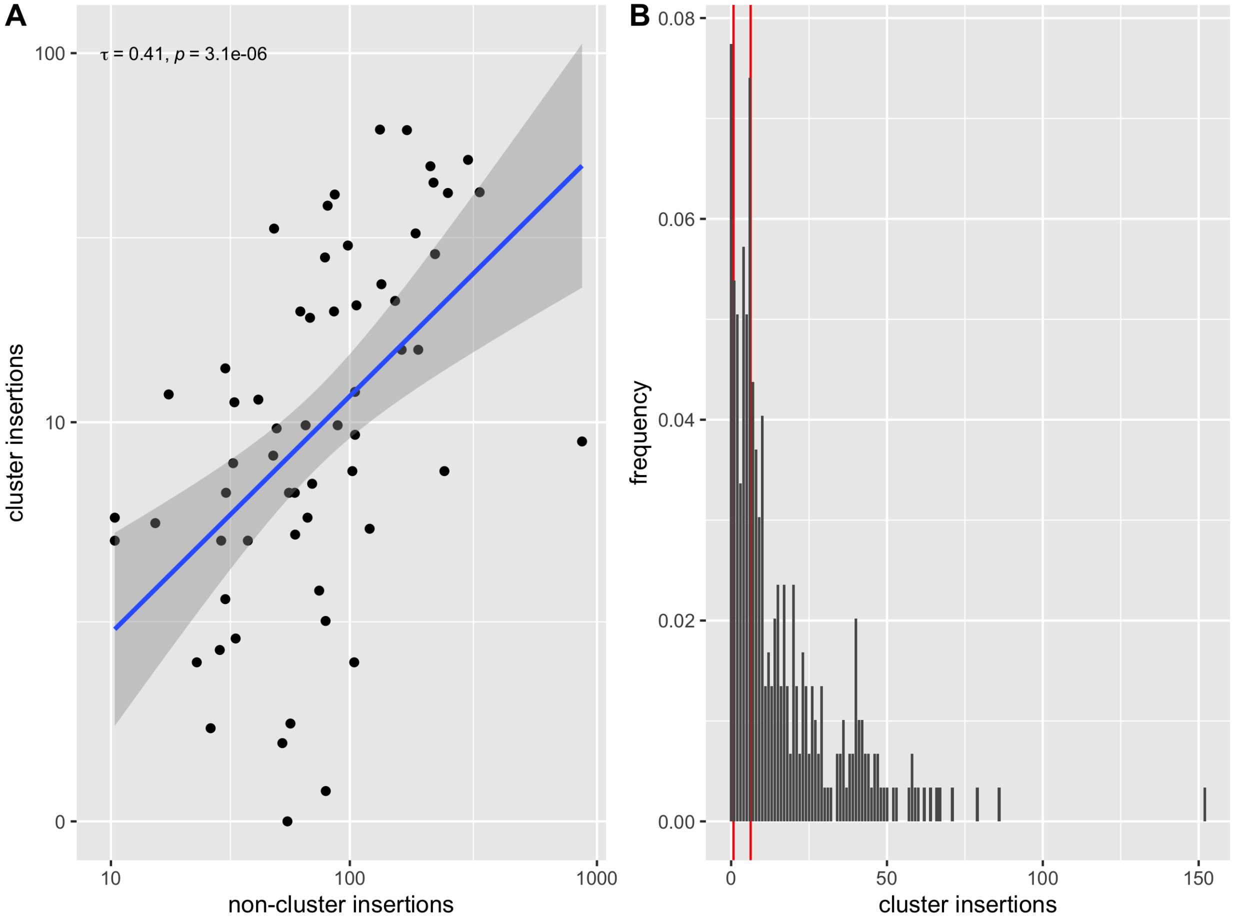

Figure S9: Correlation between cluster and non-cluster insertions (A) and the distribution of TE insertions in piRNA clusters (B) when solely completely assembled piRNA clusters (without assembly gap) are considered.

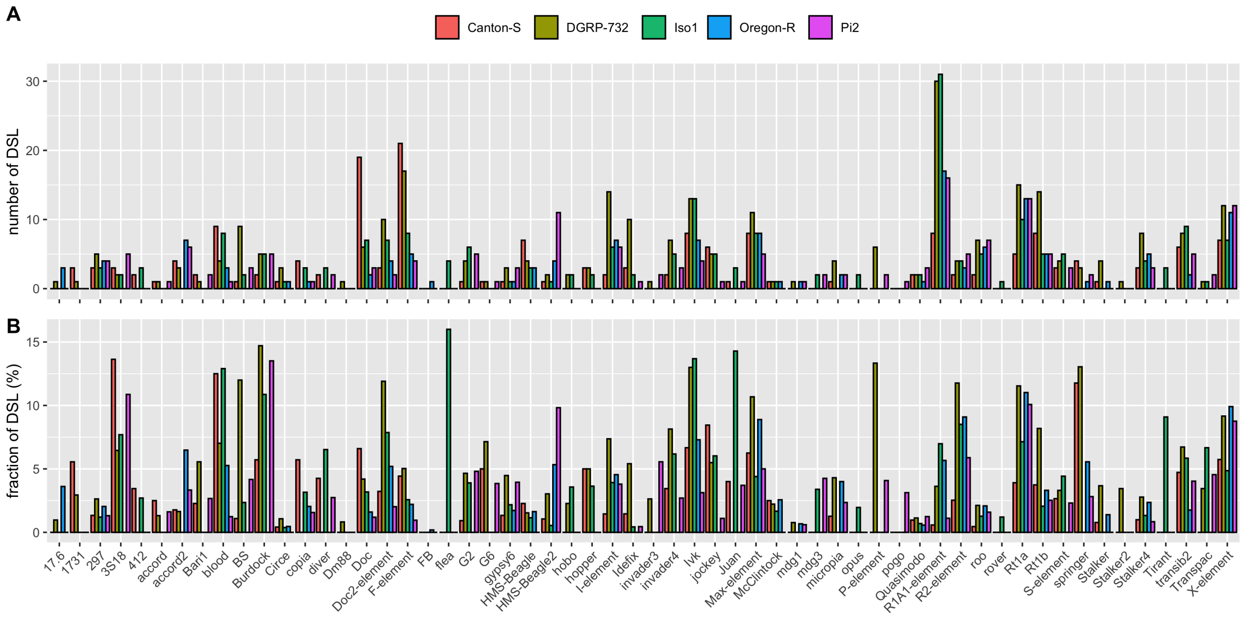

Figure S10: Heterogeneity of the abundance of DSL among TE families and fly strains. A) Absolute numbers of DSL for each TE family in the five strains. B) Fracton of DSL for each TE family in the five strains.

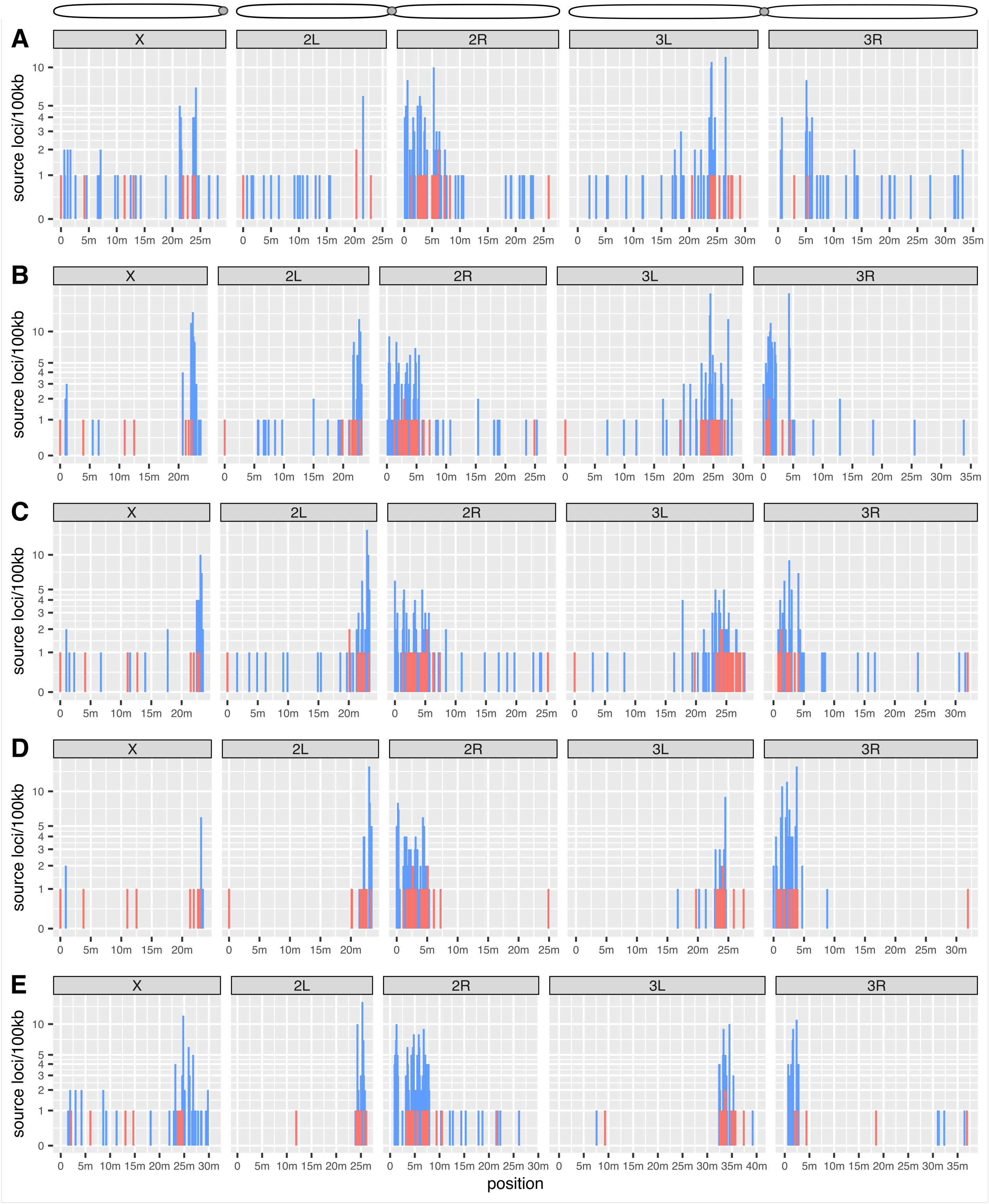

Figure S11: Distribution of piRNA source loci in the five strains (A-E). The abundance in 100kb windows is shown for TE insertions in piRNA clusters (red) and DSL (blue). The reverse complement is shown for chromosome arm 2L of Pi2 (the strand of the published assembly is likely not identical to the strand of the reference genome). A: Canton-S, B: DGRP-732, C: Iso1, D: Oregon-R, E: Pi2. A cartoon of chromosomes with gray circles corresponding to centromeres is shown above.

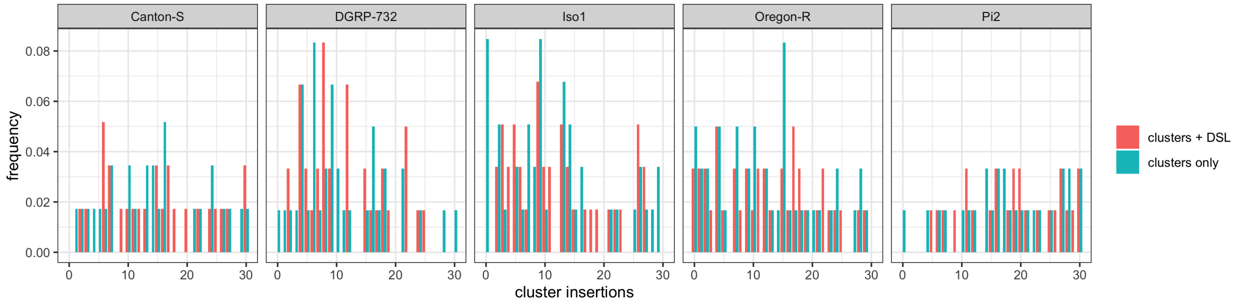

Figure S12: Abundance of piRNA producing loci in the five strains. Data are shown for cluster insertions (red) and cluster insertions plus DSL (cyan). Note that the number of families without piRNA producing locus is dramatically reduced when DSL are considered.

Table S1: General assembly statistics

| assembly | Canton-S | DGRP-732 | Iso1 | Oregon-R | Pi2 |
|---|---|---|---|---|---|
| assembly-size (MB) | 149.1 | 141.6 | 143.7 | 136.3 | 167.8 |
| N50 (MB) | 28.2 | 25.7 | 25.3 | 25.1 | 30.9 |
| BUSCO (%) | 99.4 | 99.0 | 99.5 | 99.2 | 99.3 |
| g.CUSCO (%) | 95.3 | 92.9 | 97.6 | 91.8 | 97.6 |
| u.CUSCO (%) | 78.8 | 75.3 | 85.9 | 76.5 | 81.2 |
| CQ-all | 0.069 | 0.107 | 0.141 | 0.139 | 0.066 |
| CQ-ungapped | 0.141 | 0.120 | 0.192 | 0.194 | 0.099 |
| ScQ-all | 0.512 | 0.145 | 0.099 | 0.329 | 0.351 |
| ScQ-ungapped | 0.395 | 0.162 | 0.193 | 0.317 | 0.282 |

Table S2: CUSCO of different genome assemblies. Assemblies are sorted by the u.CUSCO. Note the assemblies used in this work (bold) are among those with the highest CUSCO values.

| assembly | u.CUSCO (%) | g.CUSCO (%) |
|---|---|---|
| **Iso1** | **85.88** | **97.65** |
| GCA_020141655 | 84.71 | 97.65 |
| GCA_020142105 | 83.53 | 96.47 |
| GCA_020141675 | 82.35 | 96.47 |
| **Pi2** | **81.18** | **97.65** |
| GCA_020141955 | 80.00 | 96.47 |
| GCA_020141575 | 78.82 | 96.47 |
| **Canton-S** | **78.82** | **95.29** |
| GCA_020142005 | 77.65 | 94.12 |
| GCA_020141765 | 77.65 | 92.94 |
| **Oregon-R** | **76.47** | **91.76** |
| GCA_020141925 | 76.47 | 96.47 |
| **DGRP-732** | **75.29** | **92.94** |
| GCA_020169495 | 72.94 | 92.94 |
| GCA_020142085 | 72.94 | 96.47 |
| GCA_020141495 | 71.76 | 96.47 |
| GCA_020141935 | 70.59 | 98.82 |
| GCA_020141835 | 70.59 | 95.29 |
| GCA_020141845 | 69.41 | 92.94 |
| GCA_020141585 | 69.41 | 95.29 |
| GCA_020141795 | 65.88 | 92.94 |
| GCA_020141875 | 63.53 | 96.47 |
| GCA_020141515 | 62.35 | 85.88 |
| GCA_020141505 | 61.18 | 94.12 |
| GCA_020141595 | 58.82 | 97.65 |
| GCA_020141705 | 57.65 | 92.94 |
| GCA_020141485 | 57.65 | 92.94 |
| GCA_020142025 | 56.47 | 92.94 |
| GCA_020142045 | 55.29 | 96.47 |
| GCA_020141855 | 52.94 | 96.47 |
| GCA_020141745 | 51.76 | 94.12 |
| GCA_020141625 | 51.76 | 90.59 |
| GCA_020141815 | 49.41 | 90.59 |
| GCA_020141735 | 48.24 | 82.35 |
| GCA_020141985 | 45.88 | 87.06 |
| GCA_020141665 | 43.53 | 88.24 |
| GCA_020142055 | 36.47 | 92.94 |

Table S3: TE families without piRNA cluster insertions

| family | non-cluster | DSL | strain |
|---|---|---|---|
| R2-element | 34 | 4 | DGRP-732 |
| diver | 46 | 3 | Iso1 |
| R2-element | 47 | 4 | Iso1 |
| flea | 25 | 4 | Iso1 |
| jockey | 83 | 5 | Iso1 |
| Tirant | 33 | 3 | Iso1 |
| Bari1 | 27 | 0 | Oregon-R |
| Tirant | 5 | 0 | Oregon-R |
| R2-element | 33 | 3 | Oregon-R |
| R2-element | 85 | 5 | Pi2 |

Table S4: TE families considered for analyses in this work. TE families are sorted by their average population frequency based on Kofler et al. [2015].

| TE family | seqID | order | population frequency (%) |
|---|---|---|---|
| P-element | PPI251 | TIR | 2.9 |
| Tirant | TIRANT | LTR | 3.2 |
| R2-element | DMRER2DM | non-LTR | 3.4 |
| Stalker | STALKER | LTR | 3.5 |
| blood | BLOOD | LTR | 3.9 |
| copia | DMCOPIA | LTR | 3.9 |
| mdg1 | DMRTMGD1 | LTR | 3.9 |
| 412 | 412 | LTR | 4 |
| rover | ROVER | LTR | 4 |
| springer | SPRINGER | LTR | 4.2 |
| GATE | DME010298 | LTR | 4.3 |
| Stalker2 | STALKER2 | LTR | 4.3 |
| Transpac | AF222049 | LTR | 4.4 |
| mdg3 | DMMDG3 | LTR | 4.4 |
| jockey | DMLINEJA | non-LTR | 4.6 |
| opus | OPUS | LTR | 4.6 |
| Circe | CIRC | LTR | 4.7 |
| Doc | DMW1DOC | non-LTR | 4.8 |
| Burdock | DMU89994 | LTR | 5 |
| Max-element | DME487856 | LTR | 5 |
| R1A1-element | DMRER1DM | non-LTR | 5 |
| diver | Tinker | LTR | 5 |
| gypsy6 | GYPSY6 | LTR | 5 |
| Juan | JUAN | non-LTR | 5.1 |
| invader6 | INVADER6 | LTR | 5.3 |
| F-element | F | non-LTR | 5.5 |
| pogo | DMPOGOR11 | TIR | 5.6 |
| HMS-Beagle | Beagle | LTR | 6 |
| Dm88 | DM88 | LTR | 6.5 |
| G2 | G2 | non-LTR | 6.5 |
| McClintock | McCLINTOCK | LTR | 6.5 |
| hopper | DMTRDNA | TIR | 6.6 |
| roo | DM_ROO | LTR | 6.6 |
| NOF | FB | TIR | 6.8 |
| hobo | DMHFL1 | TIR | 6.9 |
| 3S18 | DM23420 | LTR | 7 |
| flea | DMBLPP | LTR | 7.1 |
| HMS-Beagle2 | Beagle2 | LTR | 7.2 |
| I-element | DMIFACA | non-LTR | 7.8 |
| 17.6 | DMIS176 | LTR | 7.9 |
| accord | ACCORD | LTR | 8 |
| 297 | DMIS297 | LTR | 8.1 |
| Quasimodo | QUASIMODO | LTR | 8.1 |
| Bari1 | DMBARI1 | TIR | 8.9 |
| Ivk | IVK | non-LTR | 8.9 |
| accord2 | QBERT | LTR | 9.2 |
| Rt1a | DME278684 | non-LTR | 9.7 |
| Idefix | DME9736 | LTR | 9.8 |
| Stalker4 | STALKER4 | LTR | 10 |
| FB | DMTNFB | Foldback | 10.2 |
| Rt1b | RT1B | non-LTR | 10.3 |
| BS | BS | non-LTR | 11.6 |
| invader4 | INVADER4 | LTR | 13.3 |
| G6 | G6_DM | non-LTR | 13.4 |
| invader3 | INVADER3 | LTR | 16.4 |
| X-element | ROXELEMENT | non-LTR | 17 |
| micropia | DMDM11 | LTR | 18.1 |
| Doc2-element | DOC2 | non-LTR | 20.4 |
| transib2 | TRANSIB2 | TIR | 21.1 |
| 1731 | DMTN1731 | LTR | 23 |
| S-element | DM33463 | TIR | 23.2 |
